# Quantification of biologically and chemically bound phosphorus in activated sludge from full-scale plants with biological P-removal

**DOI:** 10.1101/2021.01.04.425262

**Authors:** Francesca Petriglieri, Jette F. Petersen, Miriam Peces, Marta Nierychlo, Kamilla Hansen, Cecilie E. Baastrand, Ulla Gro Nielsen, Kasper Reitzel, Per Halkjær Nielsen

## Abstract

Large amounts of phosphorus (P) are present in activated sludge from municipal wastewater treatment plants, where it exists in the form of metal salt precipitates or biologically bound into the biomass as nucleic acids, cell membrane components, and the extracellular polymeric substances or, in special polyphosphate-accumulating organisms (PAOs), as intracellular polyphosphate. Only recently, methods that reliably allow an absolute quantification of the different P-fractions, such as sequential extraction, Raman microspectroscopy, solid-state ^31^P magic angle spinning (MAS) NMR, and solution state ^31^P NMR have been developed. This study combines these techniques to obtain a comprehensive P mass-balance of activated sludge from four wastewater treatment plants with enhanced biological phosphate removal (EBPR). The total content of P and various cations was measured by chemical analysis (ICP-OES), and different P fractions were extracted for chemical characterization. Chemically bound P constituted 38-69% of total P, most likely in the form of Fe, Mg, or Al minerals, while organically bound P constituted 7-9%. By using Raman microspectroscopy and solution state ^31^P NMR and ^31^P MAS NMR spectroscopy before and after anaerobic P-release experiments, poly-P was quantified and constituted 22-54% of total P in the activated sludges and was found in approx. 25% of all bacterial cells. Moreover, Raman microspectroscopy in combination with fluorescence *in situ* hybridization (FISH) was used to quantify the species-specific intracellular poly-P of known PAO genera (*Tetrasphaera*, *Ca.* Accumulibacter, *Dechloromonas*) and other microorganisms known to possess high level of poly-P, such as the filamentous *Ca.* Microthrix. They were all abundant, as measured by quantitative-FISH and amplicon sequencing, and accumulated large amount of poly-P, depending on their cell-size, contributing substantially to the P-removal. Interestingly, in all four EBPR plants investigated, only 1-13% of total poly-P was stored by unidentified PAO, highlighting that most PAOs in the full-scale EBPR plants investigated are now known.

**Highlights:**

- Exhaustive P mass-balance of main organic and inorganic P-species in four EBPR plants
- Quantification of poly-P of FISH-defined PAO and other species with high P content
- Total P content was 36-50 mgP/gSS of which 31-62% was in biomass and as poly-P
- A high fraction of all cells (25-30%) contained a high content of poly-P
- Known PAOs contained almost all poly-P in the EBPR plants investigated

## Introduction

Phosphorus (P) is a vital nutrient for all living organisms, but its cycle has been largely altered by human activities, resulting in increasing concerns about its supplies (Liu et al., 2016). Moreover, an uncontrolled P discharge into water bodies may cause eutrophication, which is harmful to animals and humans, but this can be partially prevented with efficient removal of P from wastewater (Oehmen et al., 2007). Nowadays, this is often achieved by introduction of the enhanced biological phosphorus removal (EBPR) process, without the need for chemical precipitants (Nielsen et al., 2019). It offers a sustainable method to recover P, where surplus activated sludge can be added to digesters and produce an effluent with a high concentration of soluble P well suited for subsequent P-recovery, e.g., as struvite, that can be used as fertilizer (Ek et al., 2006). High concentration of P is not unusual in wastewater, and in several studies it has been reported within the range of 5-15 mgP/L (Adam et al., 2009; Mino et al., 1984; Shiba and Ntuli, 2017; Wilfert et al., 2015). Worldwide, around 1.3 Mt P are treated every year in municipal wastewater treatment plants (WWTPs), which therefore represent an important source for P recovery and may theoretically substitute 40-50% of phosphate fertilizer applied in agriculture (Egle et al., 2016; Vuuren et al., 2010). Together with the EBPR process, chemical phosphorus removal (CPR) using iron or aluminum salts is the most widely applied method to remove P from wastewater. Iron salts, in particular, are extensively used because of their lower price and efficiency for P precipitation, odour control, and corrosion control (Wilfert et al., 2015, 2016).

Activated sludge typically contains 5-15 mgP/gSS for non-EBPR sludge and 30-50 mg P/gSS for EBPR sludge (Jensen et al., 2015; Menar and Jenkins, 1970; Metcalf & Eddy Inc. et al., 2014), but the P content can vary substantially among different treatment plants, comprising in general two main fractions: chemically- and biologically-bound (biogenic) P (Mino et al., 1984). In most of the cases, large amount of P is present as inorganic P, in the form of insoluble metal salts of calcium, iron, or aluminum, naturally present in the sludge, sometimes in large amounts, e.g., as vivianite (Mino et al., 1984; Wilfert et al., 2016). This fraction has been estimated in different studies and can vary from 55% to 85% of the total P (González et al., 2005; Xie et al., 2011).

Although inorganic P may be predominant in some activated sludge systems, biogenic P also plays a key role in the biomass. It is present in DNA and other nucleic acids, in cell membranes, in the extracellular polymeric substances (EPS), or in the form of orthophosphate, pyrophosphate, and polyphosphate (poly-P), normally accounting for 6.6-10.5% of total P (Mino et al., 1984; Zhang et al., 2013a). In particular, in EBPR sludge, a large fraction of biogenic P is constituted by poly-P, an intracellular storage compound typical of polyphosphate-accumulating organisms (PAO) (Nielsen et al., 2019; Oehmen et al., 2007). The EBPR process specifically selects for these microorganisms by alternating anaerobic feast and aerobic famine conditions. In the anaerobic phase, PAO take up carbon sources (e.g., acetate) and store them intracellularly as polyhydroxyalkanoates (PHA) using poly-P and, if present, glycogen hydrolysis as a source of energy. In the following aerobic phase, the stored PHA is degraded to supply growth and poly-P replenishment (Oehmen et al., 2007). Only recently, with the development of new methods, the poly-P fraction in some EBPR sludges have been quantified and reported to span between 20 and 60% of total P (Boisen et al., 2019; Fernando et al., 2019).

Activated sludge in the EBPR process comprises thousands of bacterial species, but only few genera have been recognized as PAOs. Among these are members of the genera *Ca.* Accumulibacter and *Tetrasphaera* (Nielsen et al., 2019) the most known. Both genera are common and abundant in EBPR plants worldwide (Nielsen et al., 2019), and their dynamics of storage polymers and significant contribution to P removal have been demonstrated *in situ* in several Danish WWTPs (Fernando et al., 2019). More recently, the physiology of novel PAO from the genus *Dechloromonas* has been investigated *in situ* by Raman microspectroscopy, showing a similar metabolism to *Ca.* Accumulibacter, and their potential contribution to the EBPR process (Petriglieri et al., 2020). Several putative PAOs, such as *Tessaracoccus* and *Ca.* Obscuribacter are often found in lower abundance in WWTPs (Stokholm-Bjerregaard et al., 2017), but their importance remains undescribed (Nielsen et al., 2019). Besides typical PAOs, which are cycling P in aerobic/anaerobic conditions, other microorganisms, such as *Ca.* Microthrix are known to store poly-P and release it with slower dynamics (Nielsen et al., 2002; Wang et al., 2014b, 2014a). Even though P is not cycled in these organisms as in typical PAOs, they may still contribute substantially to P removal in EBPR plants (Blackall et al., 1996; Wang et al., 2014a, 2014b). Therefore, the identification of novel PAOs and the optimization of their P uptake and release in EBPR systems is of utmost interest for process development and streamlining.

P recovery strategies rely on the specific types of P actually present in activated sludge. A range of methods exist to characterize all P fractions, such as inductively coupled plasma optical emission spectrometry (ICP-OES). Recently, a new method has been developed by Staal et al. (2019) for the characterization of different P pools in activated sludge, applying sequential P fractionations, which classifies P compounds by their chemical reactivity in specific inorganic P-pools, such as Fe-P or Ca-P (Reitzel, 2005; Staal et al., 2019). The subsequent application of solution ^31^P nuclear magnetic resonance (NMR) spectroscopy, used to identify and estimate P compounds based on the chemical bond type, can then provide information on organic P forms as well as condensed inorganic phosphates (pyro-P and poly-P) (Hupfer et al., 2008; Staal et al., 2019) in the sample. In contrast, common methods used for detection and quantification of intracellular poly-P in environmental samples are not completely reliable, as they suffer from several limitations such as inefficient extraction or degradation of poly-P (Majed et al., 2012). One of the most used techniques is staining with DNA-binding dyes followed by fluorometry, which may be unsuccessful because of the failure of the dye to penetrate cell membranes or the degradation of storage compounds during alkaline extraction (Staal et al., 2019). Raman microspectroscopy in combination with fluorescence *in situ* hybridization (FISH) provides a reliable single-cell level identification and quantification of intracellular poly-P in activated sludge (Majed et al., 2012), and it was recently used to successfully quantify poly-P content and other storage compounds in PAO cells *in situ* (Fernando et al., 2019; Petriglieri et al., 2020). The combination of Raman microspectroscopy with solid-state ^31^P magic angle spinning (MAS) NMR and solution state ^31^P NMR will therefore allow a reliable identification and quantification of total poly-P in environmental samples, including intracellular poly-P in activated sludge.

In this study, we combined a range of independent techniques to establish a robust and comprehensive P mass-balance covering all main inorganic chemical and biogenic P-species in activated sludge from four full-scale EBPR plants, which were selected based on their large size and long and stable operation. Total P was determined in activated sludge by chemical analysis (ICP-OES) and the inorganic P pools were characterized by P fractionation. The biomass was quantified by fluorescence microscopy, obtaining important information on average cell number and size, and total intracellular poly-P in the microbial biomass was estimated by Raman microspectroscopy, whereas poly-P in the bulk sludge sample was identified by solid-state ^31^P MAS NMR, and solution state ^31^P NMR. The contribution of individual PAO genera to P removal was determined by measuring the levels of intracellular poly-P in FISH probe-defined PAO cells.

## Materials and Methods

### Description of the WWTPs involved

Four full-scale EBPR WWTPs (Aalborg West, Ejby-Mølle, Lynetten, Viby) were investigated and details about their design, operation, and performance are given in Table S1. Briefly, the four plants had an alternating/recirculating design, with size between 100,000 and 1,000,000 population equivalents (PE). Two plants (Aalborg West and Lynetten) had return sludge side-stream hydrolysis (SSH). The yearly temperature range was 7-20°C. The sludge residence time in the anaerobic tanks was 2-3 h for mainstream plants and 24-28 h for the SSH plants. All plants received occasional iron dosages for improving precipitation of phosphorus and enhancing flocculation. Viby had in a period before the investigation increased Fe-dosage. All plants had good and steady operation for long time, with effluent concentrations of total P of 0.1-0.5 mg P L^−1^, and always below the limit of 1.0 mg P L^−1^. Activated sludge samples were collected from the aeration tank (at the end of aeration phase for plants with alternating operation), and transported on ice to the laboratory within few hours. Total suspended solids (TSS) was measured in accordance with Standard Methods (APHA et al., 2005). Volatile suspended solids (VSS) information was provided by the plants. An overview of the study design and all the methods applied is shown in Figure 1.

**Figure 1.**
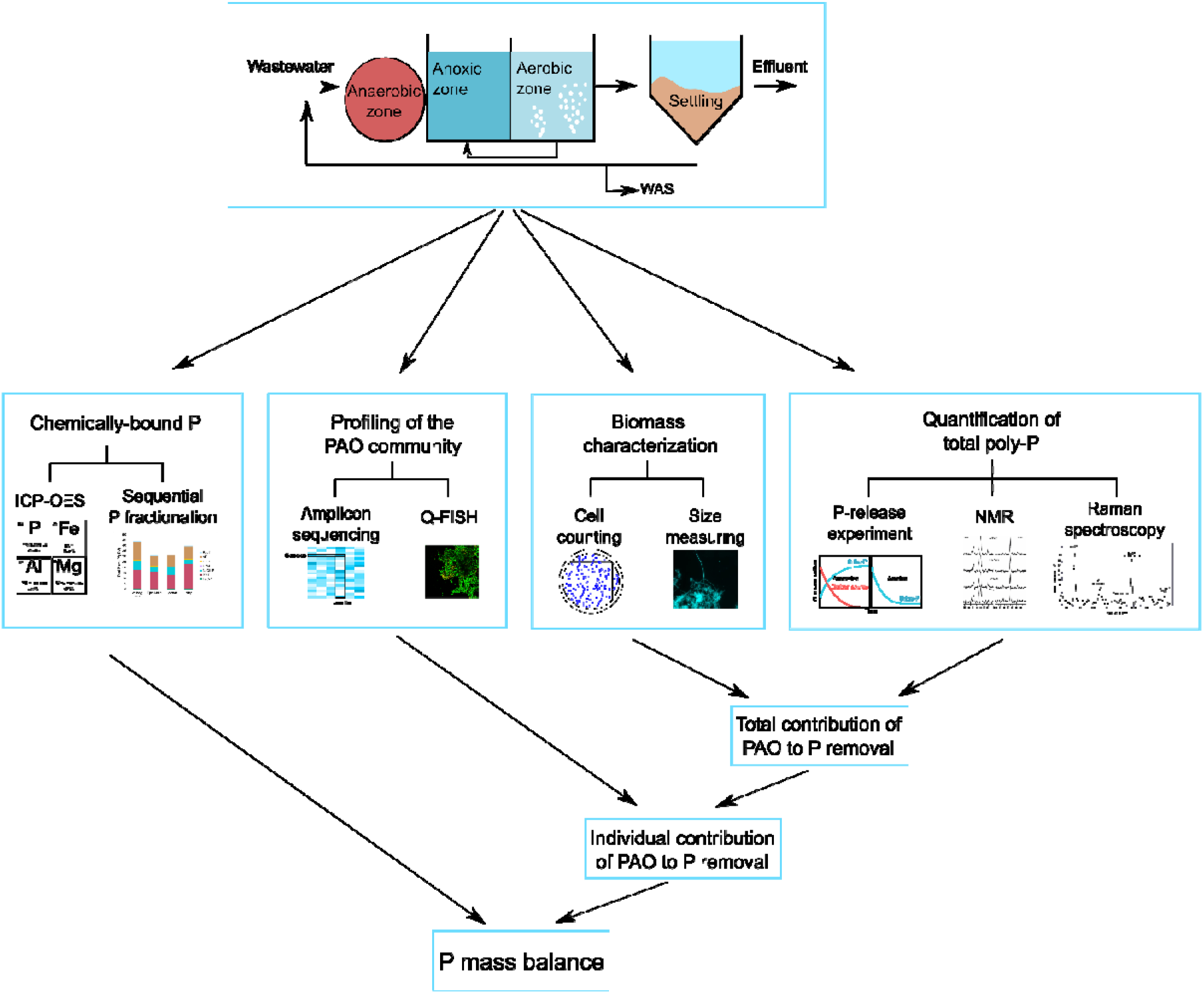
Overview of the study design. WAS = waste activated sludge; ICP-OES = inductively coupled plasma optical emission spectrometry; PAO = polyphosphate-accumulating organisms; Q-FISH = quantitative fluorescence in situ hybridization; poly-P = polyphosphate; NMR = nuclear magnetic resonance.

### Sequential P fractionation

The modified Psenner fractionation was carried out as previously described by Reitzel (2005), with a fractionation scheme that includes P associated with humic acids (Reitzel, 2005). Centrifuged activated sludge (1 g wet-weight) was used for fractionation and the scheme consisted of six sequential extraction steps: H_2_O, 0.11 M bicarbonate-dithionite (BD), 0.1 M NaOH and 0.5 M HCl, filtration and combustion of the NaOH fraction to obtain Humic-P, and a final combustion step to obtain the residual P (Res-P) as summarized in Table S2. This scheme, originally developed for lake sediments, has been previously used for investigation of P pools in activated sludge (Roske and Schonborn, 1994; Uhlmann et al., 1990). P was analyzed (as molybdate reactive P) in the 6 extracts (referred to as H_2_O-P, BD-P, NaOH-P, HCl-P, Humic-P, and Residual-P) as described above, and total P in the extracts (TP) was measured after conversion of all P in the sample to ortho-P by wet oxidation (5 mL of the fraction mixed with 1 mL potassium peroxodisulfate (K_2_S_2_O_8_, 0.2 M solution) at 120°C for 1 h). The difference between TP and ortho-P was called non-reactive P (nrP). The P fractions were normalized to the measured TP of the sludge sample, since these were associated with fewer handling steps and potential errors.

### Phosphorus release experiments from activated sludge

Batch experiments were conducted in triplicates on fresh activated sludge to analyse total P and poly-P-content per cell of FISH-defined cells under anaerobic and aerobic conditions. Fresh activated sludge samples (approx. 4 gSS/L) were aerated for 30 min to exhaust most intracellular carbon reserves and the biomass was used to measure poly-P-full cells. After aeration, sludge was transferred to 200 ml serum bottles and sealed with rubber stopper and aluminium cap. Pure nitrogen was used to flush the headspace in each bottle to ensure anaerobic conditions. A carbon source containing a mixture of acetate, glucose, and casamino acids was added to provide substrates for as many species of PAO as possible, with a final concentration of the three components of 500, 250, and 250 mg/L, respectively. The serum bottles were kept at room temperature (~22°C) with shaking for 3◻h. Samples for ortho-P analysis were collected every 20 min for the first hour of the experiment, and every 30 min during the remaining 2 hours. Biomass from the beginning (0 h) and the end of the anaerobic phase (3 h) were used to quantify poly-P and other P fractions in the sludge by solution NMR (see below). Additionally, samples from the beginning (0 h) and the end of the experiment (3 h) were flash-frozen in liquid nitrogen and subsequently freeze-dried for ^31^P MAS NMR or fixed for FISH and Raman analyses (see below).

### Chemical analyses

The ortho-P released into the liquid phase was analysed in accordance with ISO 6878:2004 using the ammonium molybdate-based colorimetric method. Total P, Fe, Al, Ca, Mg, K, and Na were measured in the fresh activated sludge samples. Sludge was homogenized for 10 sec. with a tissue homogenizer (Heidolph, Schwabach, Germany) and stored at −20°C until further analysis. Nitric acid (67%) was used to dissolve 0.5 mL of each sludge sample and the samples were microwave heated, according to (U.S. EPA 2007). The total amount of P and other elements in the samples were analysed in triplicates by ICP-OES, according to Jørgensen et al. (2017).

### Poly-P quantification by solution state ^31^P NMR spectroscopy

To identify and quantify poly-P in the sludge, solid state (see below) and solution state ^31^P NMR spectroscopy was carried out as recently described in Staal et al. (2019). Briefly, 30 mL of activated sludge were centrifuged for 10 min at 2000 rpm and the sludge pellet was used for a two-step extraction, with a 1 h pre-extraction in 40 mL of a 0.05 M EDTA solution followed by a main extraction in 40 mL of 0.25 M NaOH for 16 h. Quantitative ^31^P solution NMR spectra were recorded on a JEOL ECZ 500R 500 MHz spectrometer, using a 90° pulse (12 μs), 2.16 s acquisition time, 25 s relaxation delay, 512 scans, and proton decoupling. Spectra were processed with the MestReNova software using a 5 Hz line broadening with an exponential window function and with zero-filling to 64 K points (32 K points were recorded).

### Poly-P quantification by solid-state ^31^P MAS NMR spectroscopy

Quantitative solid-state ^31^P MAS NMR spectra were recorded on a 500 MHz (11.7 T) NMR spectrometer with a JEOL resonance ECZ 500R console and 11.7 T Oxford magnet using 3.2 mm triple resonance MAS NMR probe, single pulse excitation (45° pulse), 2-3 min relaxation delay (optimized on sample), 150 scans, and 15 kHz spinning. A synthetic sample of struvite, NH_4_MgPO_4_·6H_2_O, was used for quantitative determination of the diamagnetic P content in the samples and 85% phosphoric acid was used as a chemical shift reference (δ_iso_(^31^P) = 0 ppm). VnmrJ and MestrNova were used for data analyses. A detailed description of the protocol for data analyses was recently reported (Staal et al., 2019).

### Biomass fixation

Fresh biomass samples from batch reactors and full-scale activated sludge WWTP were either stored at −80°C for sequencing workflows or fixed for FISH with 96% ethanol or 4% PFA (final concentration), as previously described (Nielsen, 2009) and stored at −20°C until analyses.

### Cell counting and size measurement

Total number of bacterial cells, calculated in triplicates, was determined using an Axioskop epifluorescence microscope (Carl Zeiss, Oberkochen, Germany) after staining for 30 min with 50 μg/mL DAPI (4,6-diamino-2-phenylindoldihydrochlorid-dilactate). Areas of 1,500 cells for Raman microspectroscopy analysis (see below) was measured using ImageJ (Schneider et al., 2012). The biovolume of the cells was estimated by categorizing cells into perfect geometric shapes. The biovolume of filamentous bacteria was calculated considering filaments as perfect cylinders assuming an average cell length of 1 μm. Since rod-shape cells could not be classified as perfect cylinders or perfect spheroids, the biovolume of rod-shaped cells was calculated following Lee and Fuhrman et al. (1987) and Frølund et al. (1996) methodology. Briefly, the biovolume of rod-shape cells was estimated averaging the volume calculated considering rods as perfect cylinders with half-sphere ends and the volume calculated considering rods as perfect spheroids. The average width of filaments and rod-shaped cells was obtained by 2D image analysis using ImageJ (Schneider et al., 2012) for 1,000 cells, respectively.

### 16S rRNA gene amplicon sequencing

Composition of the microbial community was quantified using FISH and amplicon sequencing. DNA extraction of activated sludge samples from the MiDAS collection (McIlroy et al., 2017) was performed as described by Stokholm-Bjerregaard et al. (2017). Briefly, DNA was extracted using the FastDNA spin kit for soil (MP Biomedicals), following the manufacturer’s indications, but with an increase of the bead beating to 6 m/s for 4×40 s, using a FastPrep FP120 (MP Biomedicals). Full-length 16S rRNA gene sequencing, preparation, and taxonomic assignment of full-length 16S rRNA gene amplicon sequence variants (FL-ASVs), and amplicon sequence variant analysis (ASV) were performed as described elsewhere (Dueholm et al., 2020). Data was analyzed using R (version 3.5.2) and RStudio software (R Development Core Team, 2008).

### Fluorescence in situ hybridization and quantitative FISH

FISH was performed as described by Daims et al. (2005) using a set of specific probes: PAO651 (Crocetti et al., 2000) targeting *Ca.* Accumulibacter, MCX840 (Nierychlo et al., unpublished) targeting the genus *Ca.* Microthrix, Bet135 (Kong et al., 2007), and Dech443 (McIlroy et al., 2016) targeting the most abundant species of *Dechloromonas,* and Actino658 (Kong et al., 2005), Tetra183, and Tetra617 (Dueholm et al., 2019) to target the genus *Tetrasphaera*. Details about the optimal formamide concentration and use of competitors or helper probes can be found in Table S3. Quantitative FISH (qFISH) biovolume fractions of individual taxa were calculated as a percentage area of the total biovolume, hybridizing the EUBmix probes (Amann et al., 1990; Daims et al., 1999), that also hybridizes with the specific probe. Microscopic analysis was performed with a white light laser confocal microscope (Leica TCS SP8 X). qFISH analyses were based on 30 fields of view taken at 630× magnification using the Daime image analysis software (Daims et al., 2006).

### Raman microspectroscopy

Raman microspectroscopy was applied in combination with FISH as previously described by Fernando et al., (2019) to measure intracellular content of poly-P and other storage polymers in the cells from the beginning (0 h) and the end (3 h) of anaerobic P-release experiments. Briefly, FISH was conducted on optically polished CaF_2_ Raman windows (Crystran, UK), which give a single-sharp Raman marker at 321 cm^−1^ that serves as an internal reference point in every spectrum. Specific FISH probes (labelled with Cy3 dye) (Table S3) were used to locate the target cells for Raman analysis. After bleaching, spectra from single-cells were obtained using a Horiba LabRam HR 800 Evolution (Jobin Yvon, France) equipped with a Torus MPC 3000 (UK) 532 nm 341 mW solid-state semiconductor laser. The Raman spectrometer was calibrated prior to obtaining all measurements to the first-order Raman signal of silicon, occurring at 520.7 cm^−1^. The incident laser power density on the sample was attenuated down to 2.1 mW/μm^2^ using a set of neutral density (ND) filters. The Raman system was equipped with an in-built Olympus (model BX-41) fluorescence microscope. A 50X magnification, 0.75 numerical aperture dry objective (Olympus M Plan Achromat- Japan), with a working distance of 0.38 mm, was used throughout the work. A diffraction grating of 600 mm/groove was used, and the Raman spectra collected spanned the wavenumber region of 200 cm^−1^ to 1800 cm^−1^. The slit width of the Raman spectrometer and the confocal pinhole diameter were set respectively to 100 μm and 72 μm. Raman spectrometer operation and subsequent processing of spectra were conducted using LabSpec version 6.4 software (Horiba Scientific, France). All spectra were baseline corrected using a 6^th^ order polynomial fit.

### Determination of total poly-P and other storage polymers in all bacteria and in PAOs

The determination of the amount of intracellular poly-P and other storage polymers was carried out as described by Fernando et al. (2019). The method assumes that the intensity of the Raman signal is directly dependent on the amount of the analyte in a determined area. Using the same settings as applied in the cited paper and knowing the area of the analysed cells, it was possible to calculate the amount of poly-P and other storage polymers in chosen cells. The same principle has been applied in this study to calculate the intracellular poly-P content of the total biomass. It was measured by Raman microspectroscopy as an average of 1,500 cells randomly selected during the analysis in samples collected from the aeration tanks at the end of the aerobic phase (0 h) and after lab-scale release during anaerobic conditions (3 h) (see Supplementary text 1). An average amount of poly-P per cell was calculated as a factor of a constant determined during calibration for poly-P (Fernando et al., 2019), the average charge-coupled device (CCD) counts determined during the experiment, and the average area of cells measured by image analysis (see “Cell counting and size measuring”). This average value of poly-P per cell was multiplied by the total number of bacterial cells, counted after DAPI staining (see “Cell counting and size measuring”). As some filamentous bacteria are known to possess inclusions of storage polymers including poly-P (Nielsen et al., 2002; Wang et al., 2014b), they were included in the analysis. As it for many filaments is very difficult to determine the area of single cells inside filaments with common microscopic techniques, an arbitrary area was determined by multiplying the width of each filament measured with image analysis by 1 μm, obtaining in this way the amount of poly-P in an average segment of 1 μm length. Distribution analysis of the poly-P content and cell size was performed using R (version 3.5.2) (R Development Core Team, 2008) and RStudio software.

## Results and Discussion

The overall approach and overview of the study design is shown in Figure 1. We applied several independent methods to establish a comprehensive and robust mass balance of activated sludge from 4 full-scale EBPR plants covering P-species both chemically- and biologically-bound. These general techniques showed a high content of P (36-50 mgP/gSS), of which a substantial fraction (31-62%) was part of the biomass in the form of organic P and poly-P. The individual contribution of the known PAO to P removal indicated that most of the important PAOs in the plants investigated were described.

### Overall chemical composition of the four activated sludge

The main inorganic element besides P of the four activated sludge samples were Ca, Na, Fe, and K (Table 1). Mg, Al, and Mn were also present in lower amounts. All these elements are normally present in wastewater and may form insoluble salts with P, in particular Fe^3+^ and Al^3+^. P can be found in complexes with iron oxides or as iron phosphate minerals, such as vivianite or strengite (Wilfert et al., 2015, 2018). Ca^2+^ or Mg^2+^ can also form stable precipitates with P, and calcium phosphate precipitation in the form of apatite often takes place in EBPR systems (Manas et al., 2011; Zhang et al., 2015). Moreover, these cations are also involved in the PAO metabolism and intervene stabilizing the intracellular poly-P chains (Schonborn et al., 2001; Zhang et al., 2015). Struvite (NH_4_MgPO_4_·6H_2_O) is also a potential precipitate of P, and its accumulation in the pipes can create operational problems in the plants, causing an increase of the maintenance costs (Charles et al., 2006). However, its use for P recovery has been recently exploited, with the development of struvite precipitation from sludge or anaerobic digesters and its direct application as fertilizer (Yuan et al., 2012).

**Table 1.** Chemical composition of the four different activated sludges measured by ICP-OES. Values are given in mg/gSS.

| WWTP | Total P | Na | K | Ca | Mg | Mn | Fe | Al |
| --- | --- | --- | --- | --- | --- | --- | --- | --- |
| Aalborg West | 49.68 ± 1.89 | 74.63 ± 25.63 | 23.03 ± 0.64 | 53.83 ± 1.51 | 14.09 ± 0.43 | 0.15 ± 0.00 | 15.89 ± 0.64 | 3.40 ± 0.33 |
| Ejby Mølle | 35.85 ± 2.87 | 49.53 ± 18.12 | 18.58 ± 1.00 | 62.35 ± 8.35 | 10.18 ± 1.92 | 0.32 ± 0.12 | 27.64 ± 4.88 | 3.16 ± 0.11 |
| Lynetten | 37.10 ± 1.74 | 68.88 ± 25.45 | 18.00 ± 1.39 | 54.12 ± 5.50 | 13.44 ± 1.55 | 0.23 ± 0.12 | 25.75 ± 4.65 | 3.40 ± 0.19 |
| Viby | 45.18 ± 2.26 | 50.38 ± 22.75 | 15.28 ± 0.29 | 54.96 ± 1.36 | 8.64 ± 0.28 | 0.41 ± 0.02 | 52.24 ± 2.67 | 4.30 ± 0.43 |

### Characterization of P-pools by sequential P fractionation

To obtain an overview of the major P-pools in the four sludge samples, a sequential P fractionation was carried out (Figure 2, Table S4). Total P in the solid fraction was in the range of 31-44 mgP/gSS, and was dominated by inorganic P, extracted mainly in the BD-P and NaOH-P pools, which constituted 13-22 and 3-7 mgP/gSS, respectively, accounting together for 57-77% of the total P. The BD-P fraction is usually constituted by reducible species of Fe and Mn (Roske and Schonborn, 1994). The molar Fe:P ratio, calculated using the results obtained with ICP-OES (Table 1), was in the range of 0.31-1.15, much lower than the expected for P bound to Fe hydroxides (8.4 molar ratio) (Jensen et al., 1992) in all plants, and even lower than in common precipitates such as strengite (FePO_4_·2H_2_O) and vivianite (Fe_3_(PO_4_)_2_·8H_2_O) of 1:1 and 1.5:1, respectively (Wilfert et al., 2015, 2018), with the only exception of Aalborg West. Thus, the BD extract was not specific for extracting P bound to amorphous Fe hydroxides solely. One likely reason for this could be release of bacterial poly-P under the reducing conditions in the BD extract, as previously reported (Uhlmann et al., 1990).

**Figure 2.**
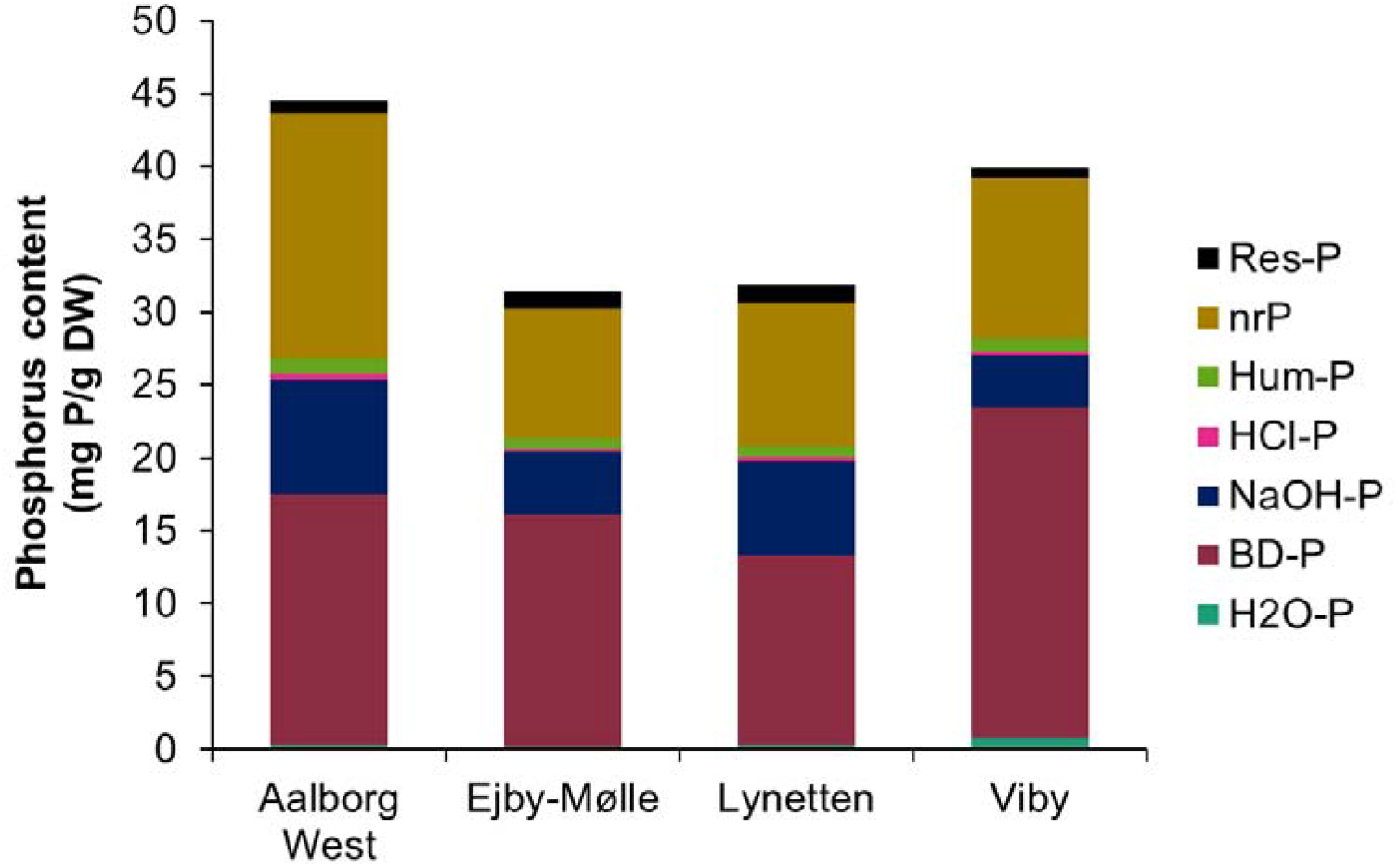
Extraction of P-fractions in activated sludge from 4 EBPR plants. The fraction H_2_O-P (water extractable P) is generally constituted by pore water P and loosely adsorbed P (<1% of total P). The BD-P (reductant soluble P) fraction is generally constituted by P bound to reducible species of Fe and Mn (inorganic fraction). The NaOH-P (NaOH soluble P) liberates P from compounds susceptible to anion exchange of PO_4_^3-^ with OH^−^ (e.g., Al oxides, Fe oxides or clay minerals). The HCl-P (HCl soluble P) fraction mainly consists of inorganic P, bound to Ca and Mg. Hum-P (Humic acid P) is constituted by P bound to humic acids. The nrP (non-reactive P) fraction is calculated by subtracting the H_2_O-P, the BD-P, and the NaOH-P from total P and contains the majority of the organic P. Res-P (residual P) is constituted by hardly degradable organic P compounds and P compounds that are not extracted in the previous steps (e.g., hematite).

Biogenic P (organic P and poly-P) was mainly extracted in the nrP pool, which accounted for 8-16 mgP/gSS (30-38% of total P). Such high nrP fraction was expected due to the abundance of microbial biomass responsible for the EBPR process, which contains poly-P, in addition to DNA-P, RNA-P, phospholipids, and pyro-P. The presence of large amounts of poly-P was confirmed by solution ^31^P NMR (see below), with similar results to previous findings (Roske and Schonborn, 1994; Uhlmann et al., 1990). Generally, the nrP quantified by ^31^P NMR spectroscopy (the sum of organic P species, pyro-P, and poly-P) was slightly higher than the nrP pool from the fractionation (Figure 2, Table S4), most likely due to extraction bias or degradation of poly-P during BD-P extraction, as previously mentioned.

### Characterization of the bacterial biomass

The bacterial biomass is an important part of the activated sludge, and we wanted to make a comprehensive P mass-balance for this fraction. We applied a microscopy-based approach, which included number of cells in the activated sludge, their sizes, and their content of poly-P (see next section). Several studies have previously attempted to quantify the number and size of bacterial cells in wastewater and activated sludge (Table 2). The preferred method is staining with fluorescent dyes that bind to the cells’ DNA, such as DAPI or acridine orange (AO), followed by epifluorescence microscopy. Different studies have reported cell numbers in the range of 2.0 × 10^11^ – 26.3 × 10^11^ cells/gVSS (Table 2) (Cloete and Africa, 1988; Frølund et al., 1996; Nielsen and Nielsen, 2002; Rasmussen et al., 1994; Urban et al., 1993; Vollertsen et al., 2001). More recently, flow cytometry has been used for rapid and direct quantification of bacterial cells in activated sludge, with similar results, in the range of 2.7 × 10^11^ – 11.0 × 10^11^ cells/gVSS (Foladori et al., 2010; Ma et al., 2013). In our study, the average number of cells obtained by DAPI staining and epifluorescence microscopy provided results in the same range of 3.78 – 6.46 × 10^11^ cells/gVSS, and very similar for all four EBPR plants (Table 2).

**Table 2.**
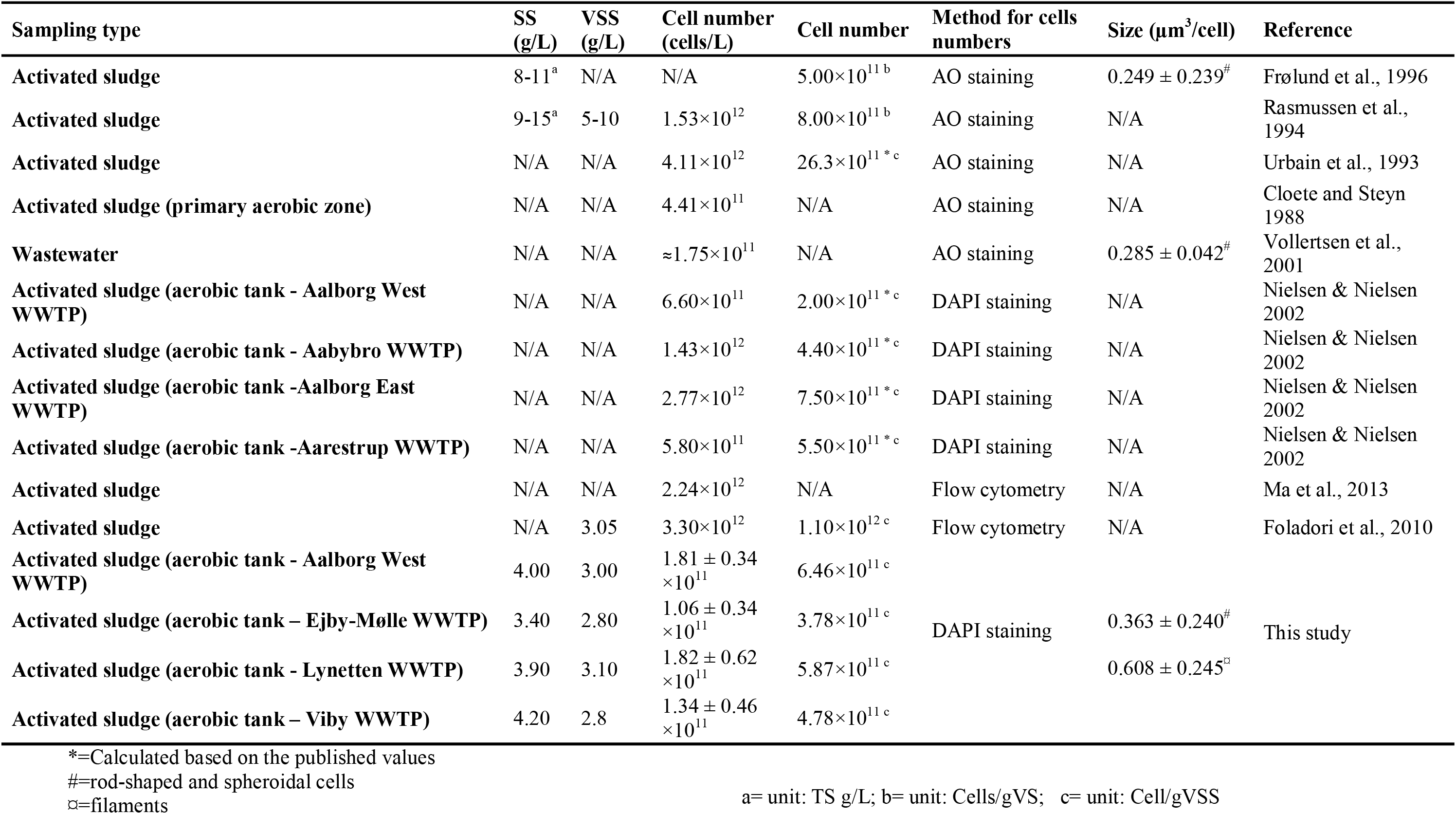
Summary of estimates of bacterial cell number and cell size in activated sludge.

The biovolume of bacteria in wastewater treatment systems has been quantified by fluorescence microscopy (Frølund et al., 1996) or flow-cytometry (Foladori et al., 2010), both providing average cell biovolume of 0.16-0.43 μm^3^. However, these estimates do not take into consideration filamentous bacteria which typically constitute 10-20% of the biomass (Nierychlo et al., 2020). Therefore, the biovolumes were measured separately for spherical/rod-shaped cells and filamentous microorganisms. In accordance with previous findings, the average biovolume of spheroidal/rod-shaped cells for the 4 plants was 0.363 ± 0.240 μm^3^, while the filaments biovolume was approximated to 0.608 ± 0.245 μm^3^ with no difference between the plants. The average length of the cells within the filaments were assumed to be 1 μm, as it is difficult to measure the real dimension of cells inside filamentous bacteria (Eikelboom, 1975; Van Veen, 1973; Ziegler et al., 1990).

### Quantification of total poly-P by different methods

Since intracellular poly-P content is regarded difficult to analyse reliably, we have applied several independent analytical tools for its absolute quantification. During anaerobic P-release experiment, active PAOs release most of their poly-P as soluble phosphate in presence of a substrate. Hence, the amount of P released during this phase was compared to the change in intracellular poly-P content obtained by Raman microspectroscopy, ^31^P MAS NMR for total poly-P in the sludge, and ^31^P solution state NMR after extraction (Table 3). All methods have recently shown to be successful in quantifying poly-P in activated sludge (Fernando et al., 2019; Staal et al., 2019a).

**Table 3.**
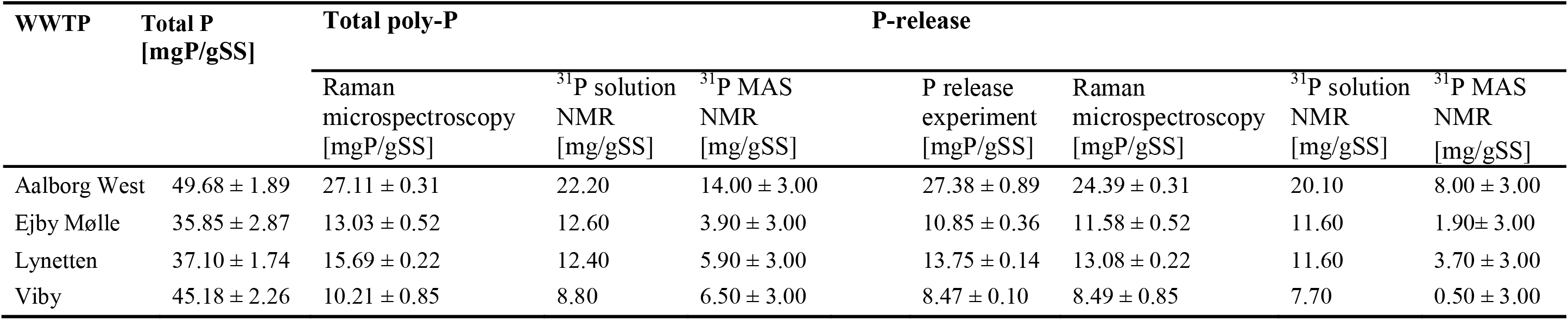
Total P, total poly-P, and anaerobic P-release in activated sludge measured by independent methods.

#### P-release test

Anaerobic P-release experiments with fresh activated sludge from the four EBPR plants were carried out to quantify the intracellular poly-P before and after anaerobic release. A mixture of different substrates, comprising acetate, glucose, and casamino acids, was used as carbon source during the experiment, to supply for a large fraction of different PAOs such as *Ca.* Accumulibacter, *Tetrasphaera*, and *Dechloromonas* (Marques et al., 2017; Petriglieri et al., 2020; Qiu et al., 2019). Similar P-release patterns were observed in all the four plants, with maximum P release after 3 h in the range of 8-13 mg P/gSS (Figure 3A, Table 3) in Lynetten, Ejby-Mølle, and Viby (Figure 3A). These values were similar to those previously obtained in Danish EBPR plants of 8-15 mg P/gSS (Mielczarek et al., 2013). The highest P release of 27.4 ± 0.9 mg/gSS was observed in Aalborg West, corresponding to the higher abundance of PAOs in this plant.

**Figure 3.**
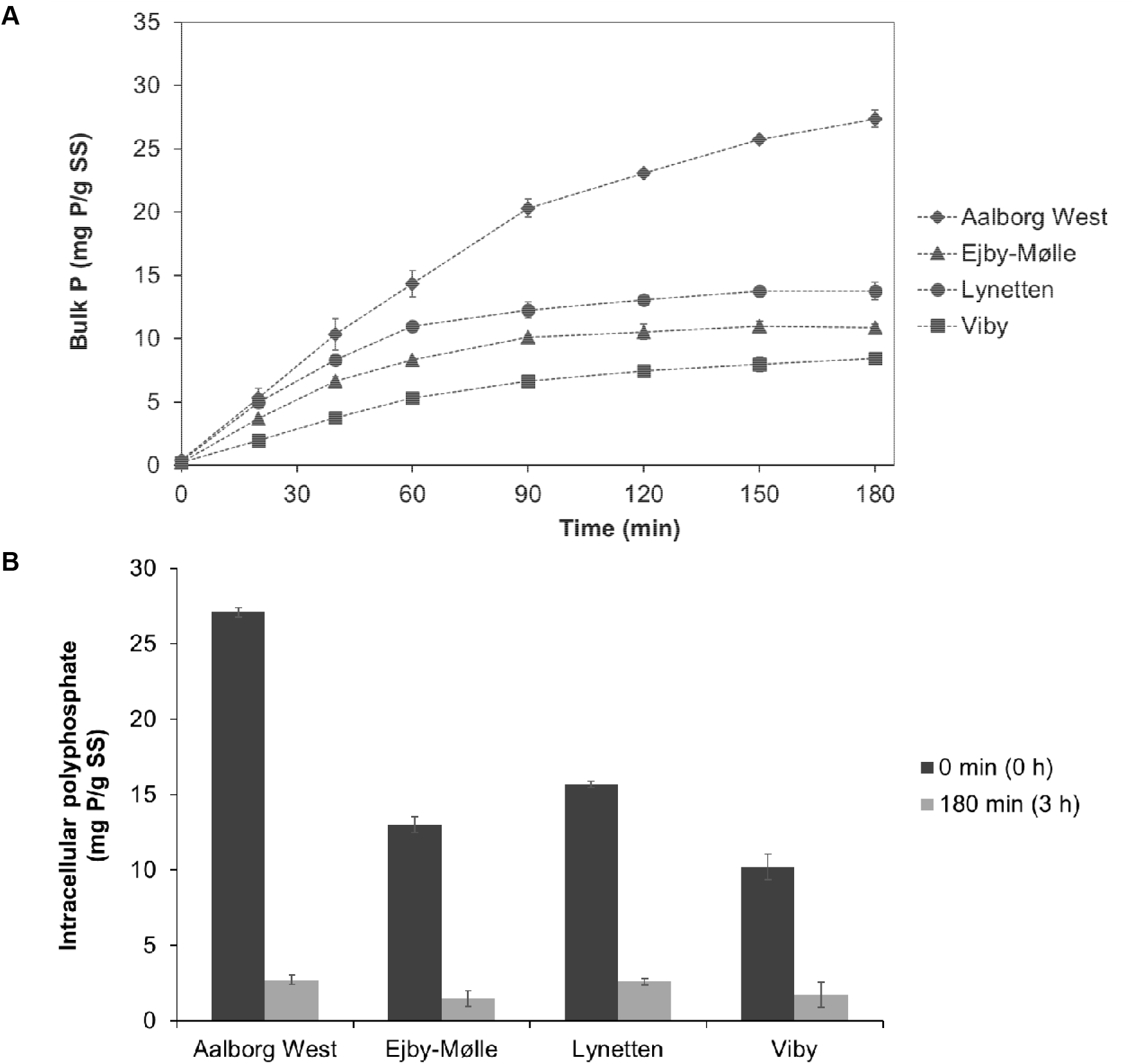
A) Bulk ortho-P concentration during the anaerobic P-release experiments with activated sludge from the four WWTPs. B) Total intracellular polyphosphate measured as an average of 1,500 randomly selected microbial cells by Raman microspectroscopy in initial samples (0 h) and after anaerobic P-release (3 h).

#### Raman microspectroscopy

Raman microspectroscopy was applied to estimate the amount of intracellular poly-P present in 1,500 randomly selected bacterial cells in the fresh activated sludge by combining Raman and DAPI staining. The total amount varied between the plants, with lowest content observed in Viby (10.2 mg/gSS) and highest in Aalborg West (27.1 mg/gSS) (Figure 3B, Table 3–4). A drastic decrease of intracellular poly-P was observed after the 3 h anaerobic conditions in sludge from all plants, reaching values of 1.72 – 2.72 mg/gSS, corresponding to 10-16% of the poly-P released (Figure 3B, Table 4). The difference in total poly-P content measured by Raman corroborated well with the P-released during anaerobic batch experiment (Figure 3A and Table 3), and with quantifications by ^31^P solution state NMR and ^31^P MAS NMR (Table 3, see below).

**Table 4.**
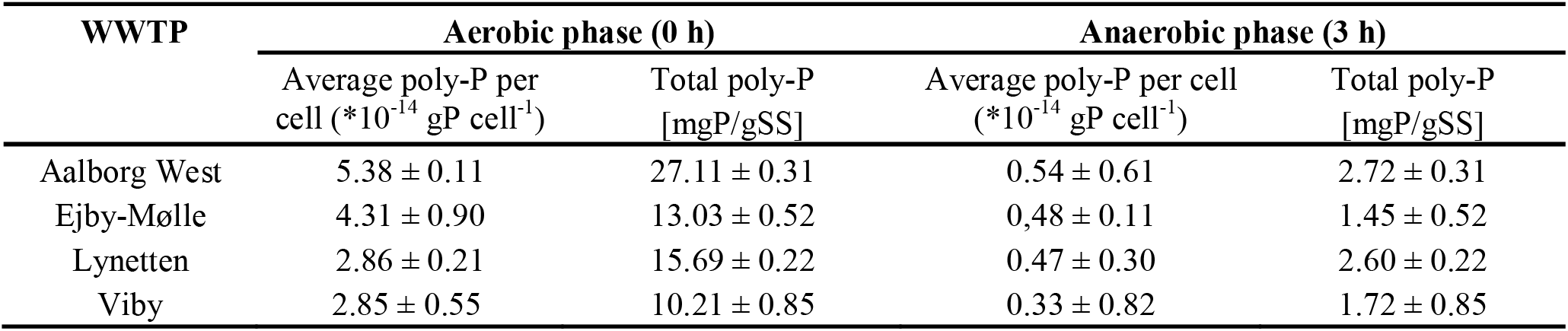
Summary of average poly-P per cell and total poly-P in activated sludge measured by Raman microspectroscopy. Total poly-P was obtained by multiplying the average poly-P per cell by total number of cells.

Interestingly, 25-32% of the bacterial cells analysed contained high amounts of poly-P (Figure 4). Among these, a fraction (5-8%) contained high amounts of poly-P also after the anaerobic incubation, thus not behaving as canonical PAOs. These may utilize other substrates or they did not show a dynamic P-cycling as observed in canonical PAOs. These could be microorganisms such as the filamentous *Ca.* Microthrix that can contain large amount of P, but do not show a typical P uptake-release dynamics (Wang et al., 2014b). Interestingly, comparison of these fractions to qFISH or amplicon sequencing results (Table 5), where the measured fraction of known PAO comprised 12-19% of the total number of bacteria, indicates that there is still up to 10% of unknown PAOs in the activated sludge samples analysed. Additionally, the lack of poly-P release in 5-10% of the cells corroborates well with the abundance of *Ca.* Microthrix, which was measured by qFISH to comprise 1.5-7.3% of the total number of bacteria. This shows that not only well-recognized PAOs such as *Ca.* Accumulibacter are important for P-removal, but also other unconventional “PAOs” such as *Ca*. Microthrix may be of key importance.

**Table 5.** Summary of relative abundances (%) of PAOs in activated sludge of the 4 WWTP measured by amplicon sequencing and q-FISH.

| WWTP | *Ca. Accumulibacter* |  | *Tetrasphaera* |  | *Dechloromonas* |  | *Ca. Microthrix* |  | Total conventional PAOs |  |
| --- | --- | --- | --- | --- | --- | --- | --- | --- | --- | --- |
|  | Sequencing | qFISH | Sequencing | qFISH | Sequencing | qFISH | Sequencing | qFISH | Sequencing | qFISH |
| **Aalborg West** | 0.9 | 3.4 ± 1.1 | 6.7 | 6.4 ± 1.8 | 5.0 | 2.6 ± 1.1 | 3.7 | 7.3 ± 2.6 | 12.6 | 12.4 |
| **Ejby-Mølle** | 0.8 | 5.2 ± 2.9 | 4.7 | 4.0 ± 1.2 | 1.5 | 1.7 ± 0.9 | 1.1 | 1.5 ± 1.1 | 7 | 10.9 |
| **Lynetten** | 0.4 | 2.6 ± 1.2 | 3.6 | 2.0 ± 0.8 | 3.6 | 5.7 ± 1.9 | 3.0 | 4.0 ± 2.3 | 7.6 | 10.3 |
| **Viby** | 0.3 | 1.0 ± 0.5 | 3.1 | 1.9 ± 0.6 | 2.8 | 2.9 ± 1.1 | 4.5 | 7.2 ± 3.8 | 6.2 | 5.8 |

**Figure 4.**
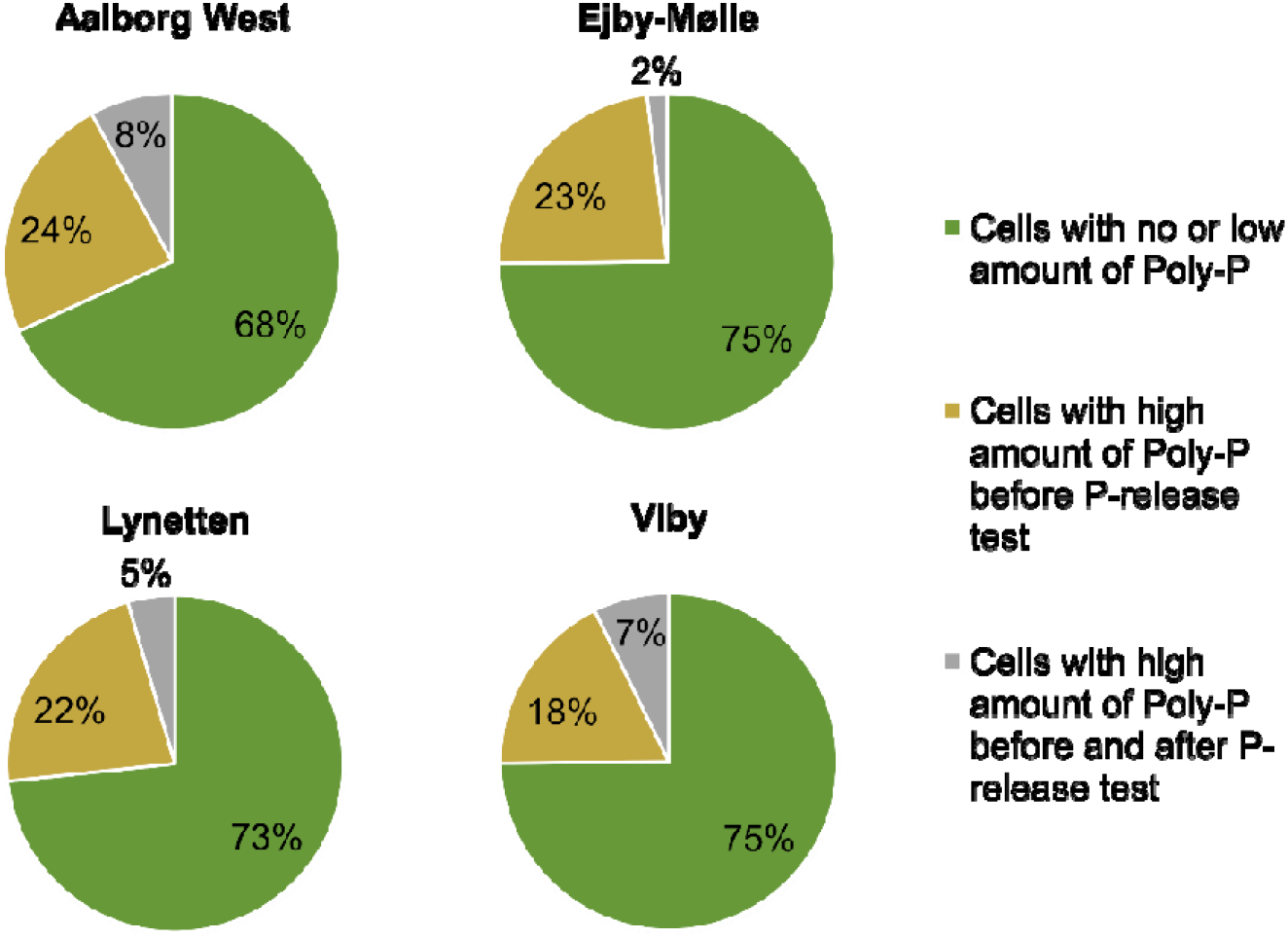
Abundance distribution of bacterial cells in activated sludge with different content of intracellular poly-P. Intracellular poly-P was measured by Raman microspectroscopy in 1,500 random DAPI stained bacterial cells before and after anaerobic P-release in the 4 full-scale plants.

#### Poly-P pools measured by ^31^P solution state NMR and ^31^P MAS NMR

Two different, but complementary, NMR-based approaches were used to calculate the total amount of poly-P in the four activated sludge samples. While ^31^P MAS NMR is used to quantify all poly-P middle groups, solution state ^31^P NMR is also showing other organic-bound P forms, as well as poly-P end groups (Staal et al., 2019). Their combined application ensures a higher accuracy in the quantification and allows the retrieval of information regarding important organic P species, such as DNA and phospholipids.

According to solution state ^31^P NMR (Table 3), poly-P constituted 8.8-22.2 mgP/gSS, which is in the similar range as previous findings of approx. 13 mgP/gSS (Staal et al., 2019), and in accordance to the amounts measured by P-release tests (8-27 mgP/gSS) and Raman microspectroscopy (10-27 mgP/gSS). In all plants, the solution state ^31^P NMR spectra of the initial samples (time = 0 h, aerobic phase) showed large peaks located at (δ(^31^P) ≈ −25 ppm originating from poly-P middle groups. There were also signals at (δ(^31^P) ≈ −5 ppm originating from poly-P end groups or pyrophosphate (Figure 5, Figures S1-S3), which would not be shown with the solely use of ^31^P MAS NMR, due to line broadening. These large peaks indicate the presence of large amounts of poly-P in the microbial biomass. In accordance with the results from the P-release experiment and the Raman analysis, the poly-P peaks were not present in the spectra of samples analysed after anaerobic release (time = 3 h) (Figure 5, Figures S1-S3), confirming the sensitivity of the methods and the dynamic levels of this poly-P in the PAO community.

**Figure 5.**
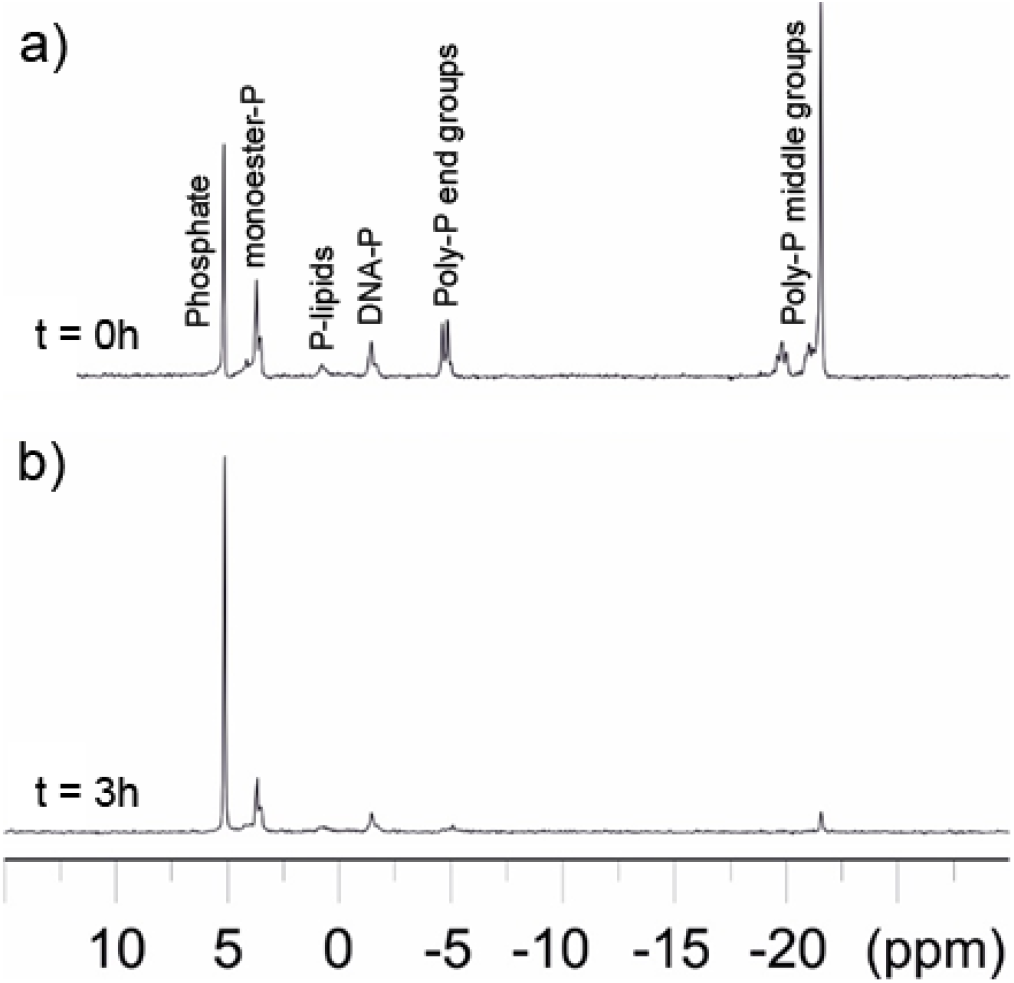
^31^P solution NMR spectra of the Ejby-Mølle samples at time 0 (a) and after 3 h anaerobic P-release test (b).

The utilization of solution state ^31^P NMR is also supplying valuable information regarding other biologically-bound P forms, such as DNA-P and P-lipids, normally present in bacterial cells as component of their membranes or genome. DNA-P was found in the biomass from all four WWTP in the range of 1.2-1.4 mg DNA-P/gSS (Table 6), which is similar to previous results found in pure cultures, in the range of 0.39 - 2.78 mg DNA-P/SS (Makarov et al., 2002). The same study also reports a total P content of 15-18 mgP/g SS for pure cultures, which is consistent with common P content in non-EBPR sludge of 5-15 mgP/g SS (Jensen et al., 2015; Menar and Jenkins, 1970; Metcalf & Eddy Inc. et al., 2014). Similarly, lipid-P values were approx. 0.5-0.6 mg DNA-P/gSS (Table 6), in the range of previously reported measurements of 0.26-9.32 for pure bacterial cultures obtained by NMR or other methods (Makarov et al., 2002; Op Den Kamp et al., 1969; Pinkart and White, 1997).

**Table 6.** Summary of total P, P content in DNA, and P content in lipids in pure cultures compared to activated sludge samples.

| Sample type | Total P<br>[mgP/gSS] | P-DNA<br>[mgP/gSS] | P-Lipid<br>[mgP/gSS] | Method for extraction and quantification | Reference |
| --- | --- | --- | --- | --- | --- |
| *Bacillus subtilis* | 15.97 – 18.38 | 0.35 – 0.39 | 8.44 – 9.32 | Extraction with 0.1 M NaOH (16 h); 31P NMR spectroscopy | Marakov et al., 2002 |
| *Bacillus subtilis* | N/A | N/A | 0.51 – 0.59 | Extraction Chloroform:Methanol:Water (1:2:0.8 v/v); Thin Layer Chromatography | Op den Kan et al., 1969 |
| *Pseudomonas putida* | 13.70 – 15.55 | 1.70 – 2.78 | 0.26 – 1.21 | Extraction with 0.1 M NaOH (16 h); 31P NMR spectroscopy | Marakov et al., 2002 |
| *Pseudomonas putida* | N/A | N/A | 2.39 – 2.61 | Extraction Chloroform:Methanol:Water (1:2:0.8 v/v); High-Pressure Liquid Chromatography | Pinkart and White 1997 |
| *Chromatiaceae* spp | N/A | N/A | 1.18 – 4.29a | Extraction with Methanol:Chloroform (2.1:0.8 v/v); Thin Layer Chromatography | Imhoff et al., 1982 |
| *Rhodospirillaceae* spp | N/A | N/A | 1.42 – 5.75a | Extraction with Methanol:Chloroform (2.1:0.8 v/v); Thin Layer Chromatography | Imhoff et al., 1982 |
| *Methylococcus capsulatus* | N/A | N/A | 3.28 | Extraction Chloroform:Methanol:Water (2:1:0.1 v/v); Thin Layer Chromatography | Makula 1978 |
| *Methylosinus trichosporium* | N/A | N/A | 3.59 | Extraction Chloroform:Methanol:Water (2:1:0.1 v/v); Thin Layer Chromatography | Makula 1978 |
| Activated sludge (Aerobic tank) |  |  |  |  |  |
| Aalborg West WWTP) | 49.7 ± 1.9 | 1.1 – 1.3 | 0.5 – 0.6 | Extraction 0.05 M EDTA (1h) + 0.25 M NaOH (16h); 31P NMR spectroscopy | This study |
| Ejby-Mølle WWTP) | 35.8 ± 2.9 | 0.9 – 1.2 | 0.4 – 0.5 | Extraction 0.05 M EDTA (1h) + 0.25 M NaOH (16h); 31P NMR spectroscopy | This study |
| Lynetten WWTP) | 37.1 ± 1.7 | 0.9 – 1.2 | 0.3 – 0.6 | Extraction 0.05 M EDTA (1h) + 0.25 M NaOH (16h); 31P NMR spectroscopy | This study |
| Viby WWTP) | 45.2 ± 2.3 | 1.1 – 1.4 | 0.1 – 0.5 | Extraction 0.05 M EDTA (1h) + 0.25 M NaOH (16h); 31P NMR spectroscopy | This study |

The ^31^P MAS NMR spectra (Figure S4) of the four initial samples (time = 0 h, aerobic phase) confirmed the results obtained by other methods, showing the spinning sideband manifold of two broad resonances located (δ(^31^P) ≈ −2 and −25 ppm, which originate from ortho- and poly-P, respectively. After 3 h, the poly-P concentration was below the detection limit for Ejby-Mølle, Viby, and Lynetten, implying full degradation of poly-P, whereas for Aalborg West ca. 9% of the total intensity was from pyro-P (Table S5). The Aalborg West sample was different from the other three plants as it had the highest concentration of P (14 mg/gSS). We note that ^31^P present in magnetic phases such as iron phosphate minerals formed e.g., by addition of Fe(III) salts to the plant, will not be observed under the experimental conditions used, as recently discussed (Staal et al., 2019). There was a small (2-4 ppm) shift in the centre of gravity of the ortho-P resonance, which contains contributions from both inorganic and organic species after 3 h, implying that the speciation of ortho-P in the sludge changed during the release experiment. At 0 h this resonance is a mixture of the different inorganic and organic orthophosphate, whereas inorganic P-phases such as magnesium and calcium P prevailed at 3 h, due to the release of biological P.

### PAO community profiling

Community profiling of the four activated sludges was performed by 16S rRNA amplicon sequencing, and FISH-based quantification was used to confirm the relative abundances obtained for target organisms (Table 5). According to the sequencing results, activated sludge in all the plants comprised a large community of PAOs, with *Tetrasphaera* being the most abundant. However, FISH (Figure 6, Table 5) showed that *Ca.* Accumulibacter and *Dechloromonas* had a larger relative biovolume (1.0 – 5.7 %) in all plants but Aalborg West, showing the difference in quantifying biomass by the two methods due to differences in extraction efficiency, primer bias, 16S rRNA gene copy number, and cell size (Albertsen et al., 2015). Furthermore, both amplicon and FISH showed that the non-conventional PAO *Ca.* Microthrix had a large relative biovolume (1.5 – 7.3%) in all plants, contributing significantly to the P-removal.

**Figure 6.**
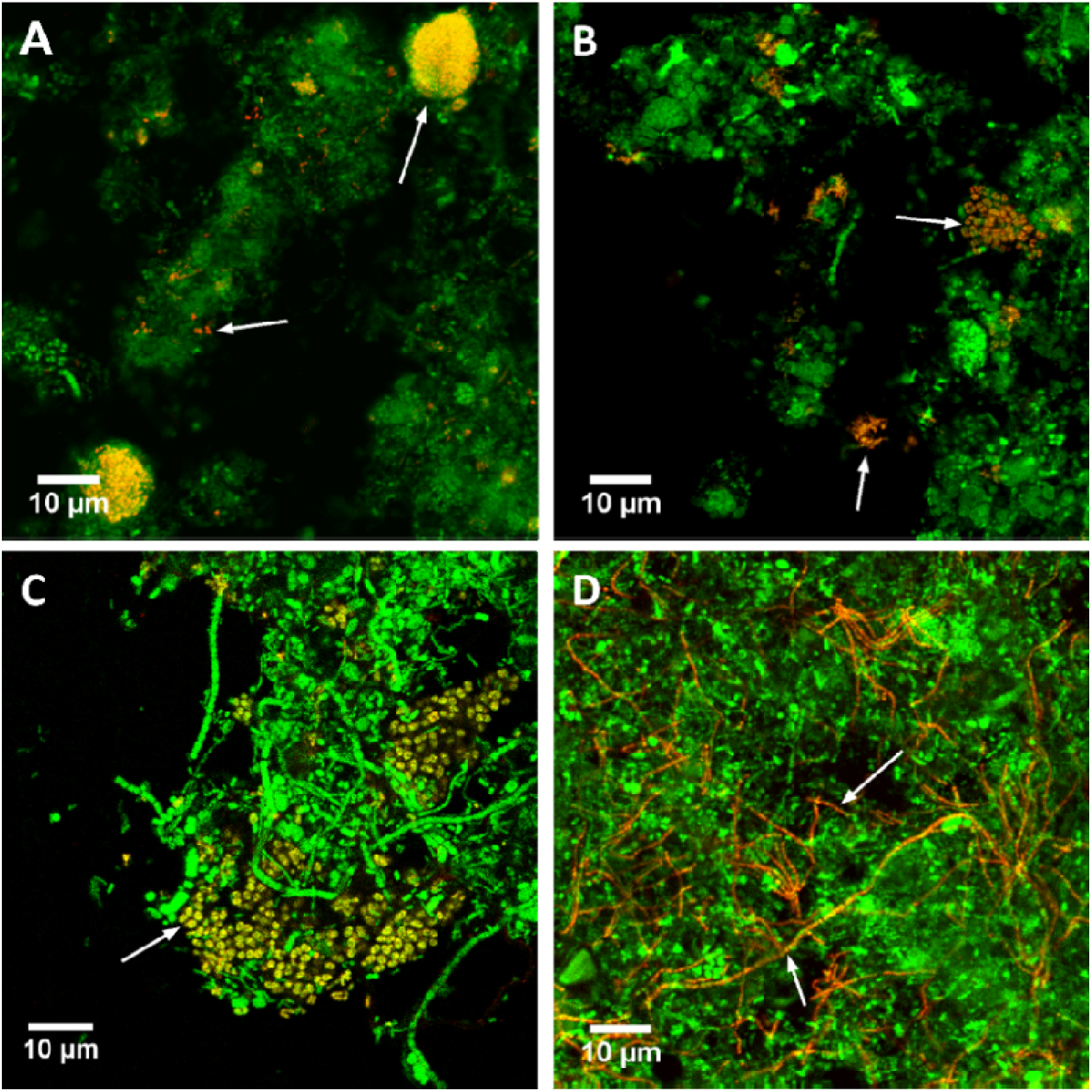
Composite FISH micrographs of PAOs in the four full-scale activated sludges. A) Specific probes (red) targeting two most abundant *Dechloromonas* species. B) Specific probes (red) targeting the *Tetrasphaera* genus. C) Specific probe (red) targeting *Ca.* Accumulibacter. D) Specific probe (red) targeting the con-conventional PAO *Ca.* Microthrix. All bacteria were targeted by EUBmix probe (green) in all images, while target bacteria appear in orange-yellow.

### Quantification of the main storage polymers in PAOs in full-scale EBPR plants

FISH-Raman was used to determine the level and dynamics of storage polymers accumulated by the known probe-defined PAOs. Moreover, the filamentous *Ca*. Microthrix, known to accumulate poly-P, but not releasing it rapidly during anaerobic conditions, was also included in the analysis. While the typical values calculated for rod-shaped cells like *Tetrasphaera* and *Ca.* Accumulibacter refer to the amount of poly-P present in a single cell, the values measured for *Ca.* Microthrix refer to a biovolume of a filament with an average 0.72 μm width and 1 μm length. The intracellular content of poly-P, PHA, and glycogen was very similar in the same probe-defined species of PAOs across different plants (Table 7-S6).

**Table 7.** Summary of the content of poly-P in probe-defined PAOs in activated sludge.

| WWTP | Poly-P in aerobic phase (0 h) (*10<sup>-14</sup> gP cell<sup>-1</sup>) |  |  |  | Poly-P in anaerobic phase (3 h) (*10<sup>-14</sup> gP cell<sup>-1</sup>) |  |  |  |
| --- | --- | --- | --- | --- | --- | --- | --- | --- |
|  | *Ca. Accumulibacter* | *Tetrasphaera* | *Dechloromonas* | *Ca. Microthrix* | *Ca. Accumulibacter* | *Tetrasphaera* | *Dechloromonas* | *Ca. Microthrix* |
| Aalborg West | 15.7 ± 0.51 | 1.68 ± 0.15 | 6.40 ± 0.23 | 1.75 ± 0.11 | 1.13 ± 0.71 | 0.49 ± 0.42 | 0.71 ± 0.36 | 1.44 ± 0.41 |
| Ejby Mølle | 14.7 ± 0.72 | 1.79 ± 0.62 | 6.95 ± 0.44 | 2.22 ± 0.71 | 1.10 ± 0.14 | 0.83 ± 0.31 | 0.86 ± 0.33 | 1.86 ± 0.62 |
| Lynetten | 15.7 ± 0.55 | 1.84 ± 0.41 | 6.52 ± 0.18 | 1.51 ± 0.22 | 1.01 ± 0.32 | 0.47 ± 0.72 | 0.65 ± 0.25 | 1.26 ± 0.31 |
| Viby | 10.7 ± 0.13 | 1.07 ± 0.71 | 6.21 ± 0.22 | 1.12 ± 0.52 | 1.46 ± 0.81 | 0.83 ± 0.61 | 0.73 ± 0.36 | 1.00 ± 0.11 |

The dynamics and levels of storage polymers in PAOs in Ejby-Mølle plant are shown as an example in Figure 6 (other plants, see Table 7–8). The amount of poly-P (g/cell) (Figure 7A) was in the range of 10-15×10^−14^, 6.2-6.9×10^−14^, 1.0-1.8×10^−14^, 1.1-2.2×10^−14^ for *Ca.* Accumulibacter, *Dechloromonas, Tetrasphaera*, and *Ca.* Microthrix, respectively. The amount calculated in this study for the well-known PAOs *Ca.* Accumulibacter and *Tetrasphaera* is lower than we measured in a previous study applying similar approach (Fernando et al., 2019).

**Figure 7.**
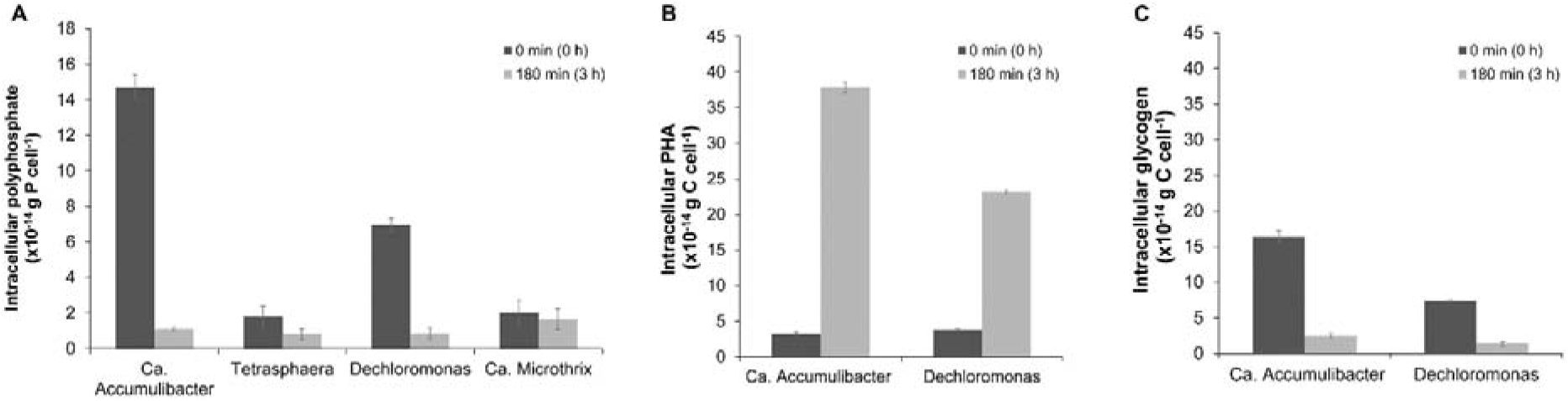
Dynamics of the storage polymers poly-P, PHA, and glycogen in important PAOs in activated sludge. Data from Ejby-Mølle WWTP is shown as example. (A) Poly-P levels during aerobic phase (0 h) and after anaerobic P-release (3 h) in known PAOs and in the unconventional PAO *Ca.* Microthrix. (B) PHA levels in *Ca.* Accumulibacter and *Dechloromonas* cells. No PHA was detected in *Tetrasphaera* and *Ca.* Microthrix. (C) Glycogen levels in *Ca.* Accumulibacter and *Dechloromonas* cells. No glycogen was detected in *Tetrasphaera* and *Ca.* Microthrix.

This difference is caused by an overestimation of the cell area by Fernando et al., (2019), as cell sizes from pure cultures or enrichments were extrapolated to full-scale plants. In this study, the area of the cells measured *in situ* were smaller, resulting in a lower amount of poly-P per cell. In all three conventional PAOs, poly-P was hydrolysed and ortho-P released into the bulk liquid during the anaerobic incubation. Considering only the poly-P amount accumulated intracellularly per cell, *Ca.* Accumulibacter had the largest amount due to its larger size (Table S7). However, when looking at the average amount of poly-P in the cells (g/μm^3^), the three conventional PAOs had almost similar content, in average 4.5×10^−14^, 3.9×10^−14^ and 2.8×10^−14^ for *Ca.* Accumulibacter, *Tetrasphaera,* and *Dechloromonas,* respectively. Moreover, in the plants investigated, all PAOs were highly abundant and, therefore, had a significant contribution to P-removal. As expected, *Ca.* Microthrix released only a small amount of poly-P during the 3 h anaerobic conditions. Although the cell width is relatively small, and thus also the P-content per μm, they can be a significant contributors to P removal with filaments length exceeding 100 μm (Blackall et al., 1996).

*Ca.* Accumulibacter and *Dechloromonas* cells possessed a dynamic behaviour of intracellular PHA and glycogen (Figure 7B-C, Table S7), as previously described (Fernando et al., 2019; Petriglieri et al., 2020) In both organisms, the C/P and C/C ratio were within the range of 0.3-0.4, in accordance with the accepted metabolic model for conventional PAOs and previous studies (Marques et al., 2017; Qiu et al., 2019). No glycogen and PHA were detected in *Tetrasphaera* and *Ca.* Microthrix, in accordance with their current metabolic models and previous findings (Fernando et al., 2019; McIlroy et al., 2013).

### A comprehensive P mass-balance

The combination of the data from ICP-OES, Raman analysis, solution state and solid state ^31^P MAS NMR, allowed us to obtain a very comprehensive and robust mass-balance of P-fractions in activated sludge from the four EBPR plants (Figure 8). Inorganic P, calculated by subtracting all the other measured fractions from the total P, was the biggest fraction in all plants except Aalborg West, constituting 38-69% of the total P. In Viby more than 70% of the total P was bound as inorganic P, most likely linked to iron or aluminium. A closer inspection of the operation in Viby revealed substantial addition of Fe(III) salts in the period before sampling, which could explain its poor EBPR performance at the time of sampling. According to the sequential P fractionations, the 57-77% of the P was in the inorganic fraction (mainly BD-P and NaOH-P fractions). The difference of these values, compared to those that can be extrapolated by solution ^31^P NMR, ^31^P MAS NMR spectroscopy, and Raman microspectroscopy measurements, were most likely due to extraction bias or degradation of poly-P during BD-P extraction.

**Figure 8.**
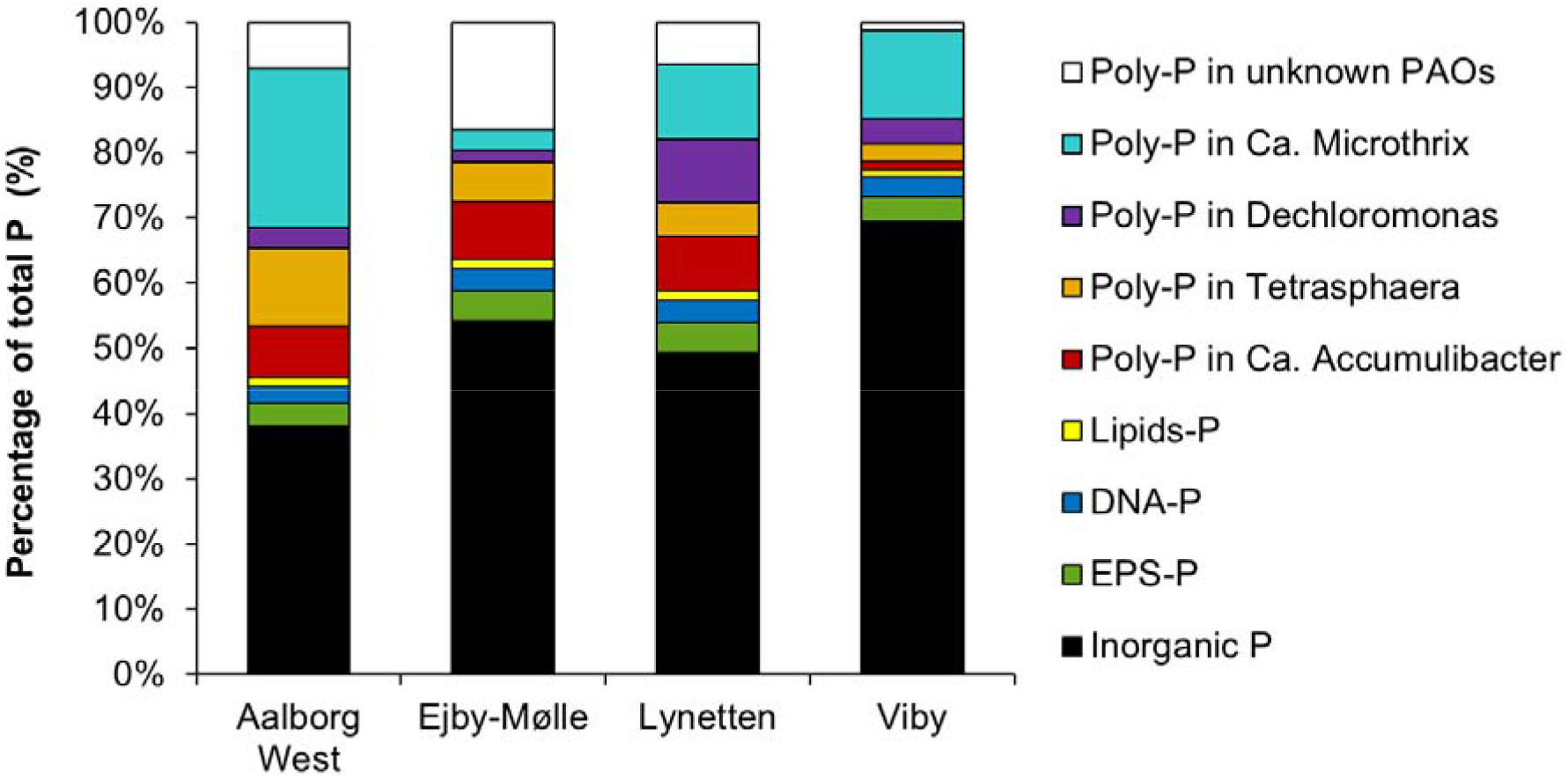
Comprehensive P-mass balance in the activated sludge. Distribution (in percentage) of P pools in activated sludge in the aeration tanks of four full-scale EBPR plants.

In all the plants, a large part of total P was present as organically-bound P, and a natural part of the microbial biomass, such as DNA-P or membrane lipids, or included into the EPS (Zhang et al., 2013a). Solution state ^31^P NMR provided valuable quantification of these distinct fractions, with DNA-P and lipid-P of approx. 1.7-1.9 mg P/gSS, constituting together 3-4% of total P (Figure 8). Previous studies on P speciation in the EPS have reported EPS-P values in the range of 0.9-2.1 mg P/gSS (Li et al., 2010; Zhang et al., 2013a, 2013b), which would account for 3-4% of the total P (Figure 8).

The second largest fraction in all plants but Viby was poly-P, constituting 22-54% of total P (Figure 8). Even though there were some variations between the independent methods applied for total poly-P quantification, all of them provided a good estimation of the poly-P content in the different samples and could therefore be independently used in further studies.

Interestingly, approx. 25% (and up to 32%) of all bacterial cells in the activated sludge accumulated large amounts of poly-P, being either PAOs or non-conventional PAOs. In all four plants, the three PAO genera investigated (*Ca.* Accumulibacter, *Tetrasphaera*, and *Dechloromonas*) were important for P-removal, and qFISH demonstrated that the known PAOs accounted for 19-20% of the biomass accumulating poly-P, leaving 12-14% undescribed. However, the results showed that most of these (2-8%) could be explained by the non-conventional PAO, *Ca.* Microthrix. Non-conventional PAOs are easily overlooked since they do not exhibit the dynamic P release/uptake in typical tests for PAO activity. In general, the contribution of the known PAO genera varied from plant to plant, depending on their relative abundances. The contribution of *Ca.* Microthrix exceeded the other known PAOs in three plants (Aalborg West, Lynetten, and Viby) because of its high abundance. This filamentous bacterium is considered unwanted because it causes settling (bulking) or foaming problems in full-scale WWTPs (Nierychlo et al., 2020), but this study also shows that they should also be regarded as important contributors to P removal in EBPR systems. Considering this non-conventional PAO, the mass-balance shows that only a small fraction of total poly-P (1-13%) was accumulated by PAOs that are still undescribed, indicating that most important PAOs are now known in the plants investigated.

### Concluding remarks

The activated sludge samples, taken from the four largest EBPR plants in Denmark, had a total P content of 36-50 mgP/gSS, and in all of them, except one, a substantial fraction of P (31-62%) was part of the biomass in the form of organic P and poly-P. The total amount of poly-P was successfully measured by several independent methods. Extensive analyses of number and sizes of all bacterial cells and specific FISH probe-defined PAOs allowed us to quantify the individual contribution of the known PAOs, *Ca.* Accumulibacter, *Tetrasphaera,* and *Dechloromonas* to P removal. In addition, the non-conventional PAO *Ca.* Microthrix contained large amounts of poly-P contributing considerably to P-removal. The distribution of poly-P in random bacterial cells showed that cells with high content of poly-P constituted 25-32% of all cells. Only a small amount of poly-P (1-13%) could not be assigned to any known PAO, indicating that most of the important PAOs in the plants investigated were described.

## Acknowledgements

We thank Susanne Bielidt for assistance with cell counting and microscopical analysis. The project was funded by the Villum Foundation (Dark Matter, grant 13351) and Innovation Fund Denmark (ReCoverP, grant 4106-00014B).

