## Supplementary Information for "Quantification of biologically and chemically bound phosphorus in activated sludge from full-scale plants with biological P-removal"

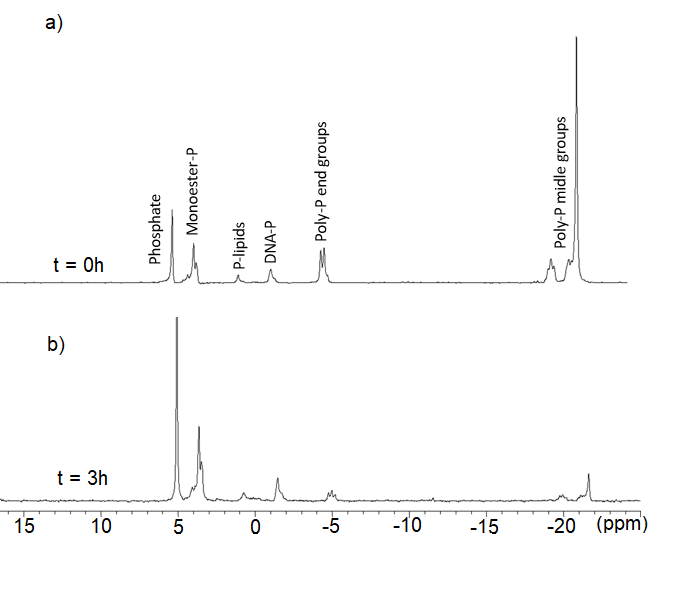

**Figure S1.** ^31^P solution NMR spectra of the Aalborg West samples at time 0 (a) and after 3 h (b).

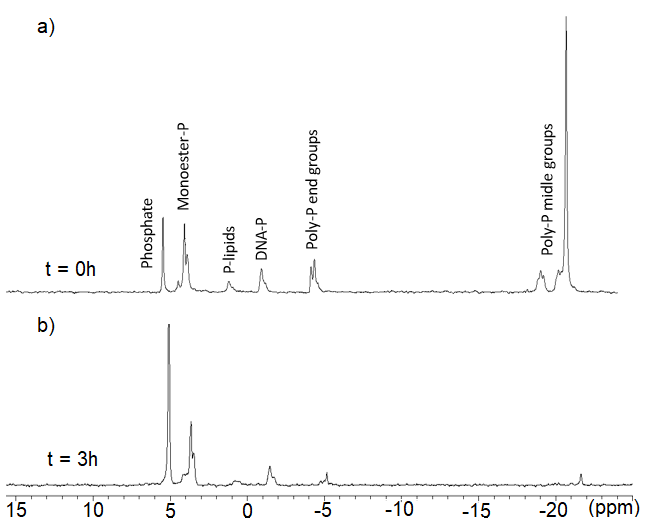

**Figure S2.** ^31^P solution NMR spectra of the Lynetten samples at time 0 (a) and after 3 h (b).

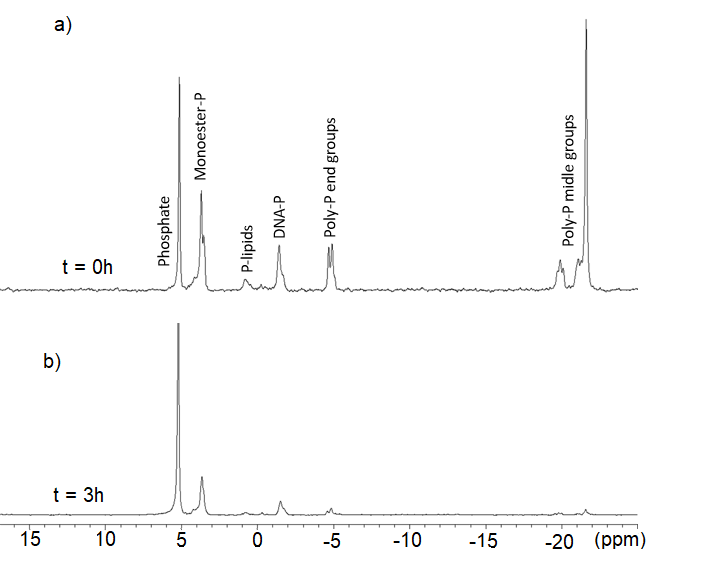

**Figure S3**. ^31^P solution NMR spectra of the Viby samples at time 0 (a) and after 3 h (b).

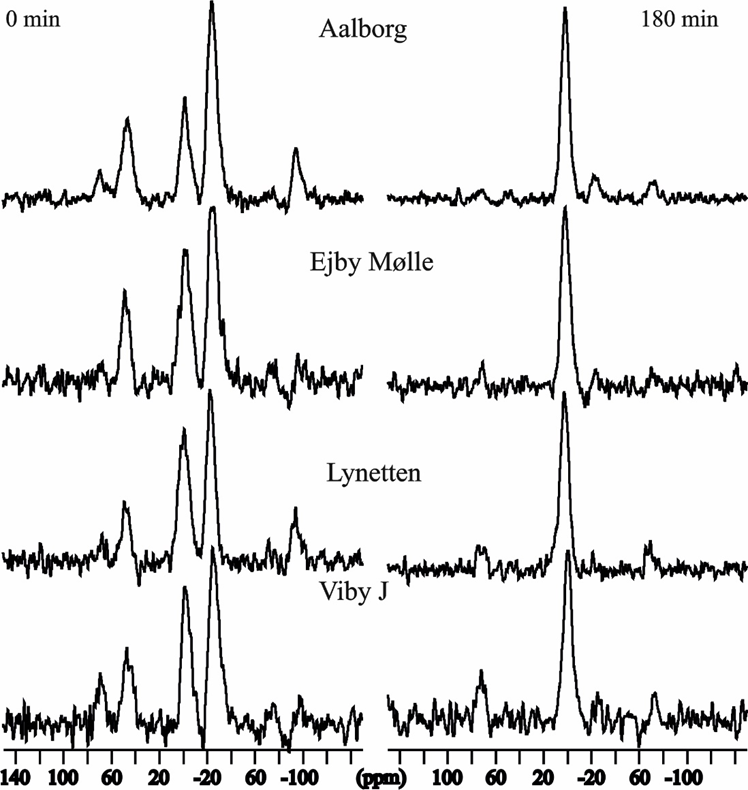

**Figure S4.** ^31^P MAS NMR spectra of the eight activated sludge samples.

**Table S1.** Details about the EBPR plants.

| **Plant** | **PE Capacity** | **Industrial Percentage** | **Operation** | **Primary settling** | **Operation (EBPR/BNR)** | **Sidestream hydrolysis** | **Anaerobic Digester Type** |
| --- | --- | --- | --- | --- | --- | --- | --- |
| Aalborg West | 330,000 | 25 | alternating | yes | EBPR | yes | thermophilic |
| Ejby Mølle | 410,000 | 49-55 | alternating | yes | EBPR | no | thermophilic |
| Lynetten | 1,000,000 | 10 | alternating | yes | EBPR | yes | mesophilic |
| Viby | 100,000 | 5 | recirculation | yes | BNR | no | mesophilic |

**Table S2.** Overview of P fractionation of P pools. Modified from Reitzel (2005).

| **Fraction** | **Extractant/procedure** | **P species extracted** |
| --- | --- | --- |
| **H2O** | Deionized H_2_O  25 mL (1 h)  25 mL (5 min) | Loosely bound  Porewater P |
| **BD** | 0.11 M Bicarbonate-dithionite  25 mL (1 h)  25 mL (5 min)  25 mL (5 min)  25 mL H_2_O (5 min)  Aeration | Reducible Fe species  Reducible Mn species  Some organic P |
| **NaOH** | 0.1 M NaOH  25 mL (16 h)  25 mL (5 min)  25 mL H_2_O (5 min) | Organic P  Anion exchangeable P  P from Al oxides  P from Fe oxides  Inorganic poly-P  Struvite |
| **nrP** | Calculated as the difference of total P and H2O-P, BD-P and NaOH-P | Organic P |
| **HCL** | 0.5 M HCl  25 mL (1 h)  25 mL (5 min)  25 mL H_2_O (5 min) | Ca-P  Mg-P  Residual organic P |
| **Residual-P** | Dried  Combusted 520°C  8 mL 1 M HCl (120°C, 1 h) | Hardly degradable organic-P  Other non-extracted compounds |
| **Humic-P** | NaOH fraction filtered  Combusted at 520°C  8 mL 1 M HCl (120°C, 1 h) | P bound to humic acids |

**Table S3.** List of FISH probes used in this study.

| **Probe** | **Target group** | **Sequence (5’-3’)** | **[FA]%^**^** | **Reference** |
| --- | --- | --- | --- | --- |
| **EUB338** | **All bacteria** | **GCTGCCTCCCGTAGGAGT** | **N/A** | (Amann et al., 1990) |
| **EUB338 II** | **All bacteria** | **GCAGCCACCCGTAGGTGT** | **N/A** | (Daims et al., 1999) |
| **EUB338 III** | **All bacteria** | **GCTGCCACCCGTAGGTGT** | **N/A** | (Daims et al., 1999) |
| **PAO651** | ***Ca*. Accumulibacter** | **CCCTCTGCCAAACTCCAG** | **35** | (Crocetti et al., 2000) |
| **Bet135** | ***Ca.* Dechloromonas phosphatis** | **ACGTTATCCCCCACTCAATGG** | **45** | (Kong et al., 2007) |
| **Dech443** | ***Ca.* Dechloromonas phosphovora** | **ACCCATGCATTTTCTTCCCGG** | **35** | (Mcilroy et al., 2016) |
| Dech443c1 | Competitor for Dech443 | ACCCATGCGTTTTCTTCCCGG | N/A | (Mcilroy et al., 2016) |
| **Actino658** | ***Tetrasphaera* midas_s_5** | **TCCGGTCTCCCCTACCAT** | **40** | (Kong et al., 2005) |
| Actino658c2 | Competitor for Actione658 | ATTCCAGTCTCCCCTACCAT | N/A | (Kong et al., 2005) |
| **Tetra183** | **Genus probe for Tetrasphaera** | **TAGAGATGCCTCTCCGTCTC** | **30** | (Dueholm et al., 2019) |
| Tetra183h1 | Helper for Tetra183 | YCCAGAGTCTGGGGCAGGTT | N/A | (Dueholm et al., 2019) |
| Tetra183h2 | Helper for Tetra183 | GTCCATCCCAGACCGAAAAACTTT | N/A | (Dueholm et al., 2019) |
| **Tetra617** | **Species not targeted by Tetra183** | **CCCACTGCAAGTCCGGAATTGAGT** | **30** | (Dueholm et al., 2019) |
| **MCX-840** | **Genus probe for *Ca.* Microthrix** | **CGGCGCGGAGAGAGTTGAGT** | **20** | (Nierychlo et al., in preparation) |
| MCX-840h1 | Helper for MCX-840 | TCTCCCCACACCTAGTGCCCAACG | N/A | (Nierychlo et al., in preparation) |
| MCX-840h2 | Helper for MCX-840 | GCGGGGCACTTAATGCGTTAGCTA | N/A | (Nierychlo et al., in preparation) |

**Table S4.** Summary of the P pools obtained by sequential fractionations and ^31^P NMR.

| **WWTP** | **H2O-P [mg/g]** | **BD-P**  **[mg/g]** | **NaOH-P [mg/g]** | **nrP**  **[mg/g]** | **Hum-P [mg/g]** | **HCl-P [mg/g]** | **Res-P [mg/g]** | **TP**  **[mg/g]** |
| --- | --- | --- | --- | --- | --- | --- | --- | --- |
| Aalborg West | 0.32±0.06 | 17.20±2.47 | 7.85±0.43 | 16.81 ±1.72 | 1.03±0.03 | 0.39±0.03 | 0.83±0.25 | 44.43±0.00 |
| Ejby Mølle | 0.20±0.01 | 15.90±1.76 | 4.32±0.70 | 8.95±0.40 | 0.72±0.03 | 0.18±0.03 | 1.12±0.05 | 31.42±0.54 |
| Lynetten | 0.32±0.03 | 13.04±0.28 | 6.41±0.23 | 9.88±0.31 | 0.72±0.17 | 0.30±0.01 | 1.23±0.07 | 32.00±0.00 |
| Viby | 0.78±0.03 | 22.68±0.25 | 3.70±0.09 | 10.99±2.65 | 0.91±0.03 | 0.13±0.01 | 0.73±0.02 | 39.92±0.24 |

**Table S5**. The absolute concentration of diamagnetic phosphate in the sample with the relative concentration of orthophosphate (%Portho) and polyphosphate (%Ppoly) determined from deconvolution of the 31P MAS NMR spectra. % indicates that the quantity is below the detection limit of the NMR experiment (< 3%). The relative uncertainty of the integrals is ca 5%.

| **Sample** | **P_nmr_ [mg/g]** | | **P_ortho_**  **%** | **P_poly_**  **%** |
| --- | --- | --- | --- | --- |
| Struvite | | 126 | 100 | - |
| Aalborg West 0 h | | 19 | 26 | 74 |
| Aalborg West 3 h | | 14 | 88 | 9 |
| Ejby Mølle 0 h | | 8.2 | 52 | 48 |
| Ejby Mølle 3 h | | 6.2 | 100 | % |
| Lynetten 0 h | | 9.7 | 39 | 61 |
| Lynetten 3 h | | 7.5 | 100 | % |
| Viby 0 h | | 11 | 39 | 61 |
| Viby 3 h | | 3.8 | 100 | % |

**Table S6.** Area and biovolume of PAO cells

| **Sample** | **Cell area (µm^2^)** | **Biovolume (µm^3^)** |
| --- | --- | --- |
| *Candidatus* Accumulibacter | 1.3 | 3.14 |
| *Tetrasphaera* | 0.38 | 0.45 |
| *Dechloromonas* | 1.10 | 2.35 |
| *Candidatus* Microthrix | 0.72 0.40  Average length: 91.4 µm | |

**Table S7.** Summary of the content of PHA and glycogen in known PAOs.

| **WWTP** | **PHA**  **(*10^-14^ gC cell^-1^)** | | | | **Glycogen**  **(*10^-14^ gC cell^-1^)** | | | |
| --- | --- | --- | --- | --- | --- | --- | --- | --- |
|  | Aerobic phase (0 h) | | Anaerobic phase (3 h) | | Aerobic phase (0 h) | | Anaerobic phase (3 h) | |
|  | *Ca.* Accumulibacter | *Dechloromonas* | *Ca.* Accumulibacter | *Dechloromonas* | *Ca.* Accumulibacter | *Dechloromonas* | *Ca.* Accumulibacter | *Dechloromonas* |
| Aalborg West | 3.40 ± 0.11 | 3.14 ± 0.28 | 39.5 ± 0.40 | 20.2 ± 0.14 | 17.7 ± 0.50 | 6.99 ± 0.17 | 2.98 ± 0.12 | 1.21 ± 0.16 |
| Ejby Mølle | 3.28 ± 0.21 | 3.79 ± 0.24 | 37.9 ± 0.70 | 23.2 ± 0.36 | 16.4 ± 0.81 | 7.37 ± 0.13 | 2.48 ± 0.31 | 1.37 ± 0.34 |
| Lynetten | 2.69 ± 0.12 | 3.22 ± 0.17 | 35.2 ± 0.31 | 22.3 ± 0.13 | 15.2 ± 0.91 | 7.61 ± 0.25 | 2.46 ± 0.60 | 1.64 ± 0.16 |
| Viby | 2.50 ± 0.51 | 3.41 ± 0.23 | 30.0 ± 0.22 | 21.3 ± 0.23 | 14.3 ± 0.72 | 7.48 ± 0.28 | 2.71 ± 0.21 | 1.55 ± 0.39 |

**Supplementary text**

1. **Example of total poly-P quantification with the Raman-based approach**

Microbial biomass in 4 different activated sludge was analyzed at the end of aerobic phase (0 h) and end of anaerobic phase (3 h), under identical instrument settings as described in (Fernando et al., 2019). Calculations for Ejby-Mølle WWTP are reported here as an example. The average Raman intensities obtained for cells in poly-P “full” and “empty” states were 87.52 CCD counts and 9.86 CCD counts respectively (n = 1500 randomly selected microbial cells). The average 2D area occupied by a single cell, when mounted on CaF2 Raman substrate was estimated to 0.99 ± 0.12 µm^2^, using the image processing software, ImageJ (n = 1,500 cells). The calibration coefficient (k) for poly-P under these conditions and instrument settings, determined in (Fernando et al., 2019), was equal to 4.98 * 10^-16^ g P μm^-2^ counts^-1^. According to the method developed by Fernando et al., (2019):

- Poly-P per cell = k × average Raman CCD counts × estimated cell area

For an average single cell at the end of the aerobic phase (poly-P “full”state):

Average poly-P per cell = 4.98 × 10^-16^ g P μm^-2^ counts^-1^ × 87.52 counts × 0.99 μm^2^/cell = 4.31× 10^-14^ g P/cell

For an average single cell at the end of the anaerobic phase (poly-P “empty” state):

Average poly-P per cell = 4.98 × 10^-16^ g P μm^-2^ counts^-1^ × 9.86 counts × 0.99 μm^2^/cell = 0.48 × 10^-14^ g P/cell

The numbers so obtained were then multiplied by the total number of cells, measured microscopically, to determine an estimation of the total poly-P:

Total poly-P (0 h) = 4.31 × 10^-14^ g P/cell × 2.7 × 10^11^cells/g SS = 11.63 mg P/g SS

Correction for poly-P loss (12%) during storage = 11.63 × 1.12 = 13.03 mg P/g SS

Total poly-P (3 h) = 0.48 × 10^-14^ g P/cell × 2.7 × 10^11^cells/g SS = 1.29 mg P/g SS

Correction for poly-P loss = 1.29 × 1.12 = 1.45 mg P/g SS
